# *Staphylococcus aureus* triggers isolate-specific host transcriptional responses alongside TNF-R1 regulated cell death

**DOI:** 10.64898/2026.05.06.723175

**Authors:** Annika Walter, Thorsten Bischler, Marvin Jungblut, Leonhard F. Breitsprecher, Johanna Beck, Lucia Hofmann, Nicole Schaefer, Tanja Ziesmann, Silke Haerteis, Iana Gadjalova, Ute Distler, Gerti Beliu, Olympia-Ekaterini Psathaki, Michael Hensel, Wulf Schneider-Brachert, Tom Graefenhan, Thomas Stempfl, Baerbel Kieninger, Sabrina Muehlen, Volker Alt, Gopala-Krishna Mannala, Juergen Fritsch

## Abstract

**Background:** *Staphylococcus aureus* (*S. aureus*) is an increasingly recognized intracellular pathogen, yet infection outcomes vary with bacterial isolate and host cell type. The mechanisms underlying these differences remain poorly understood. This study investigates how distinct intracellular *S. aureus* isolates influence host signaling programs and infection outcomes by modulating cell death pathways and TNF-R1 dependent regulation of host cell fates across different human cell lines.

**Methods:** Four *S. aureus* isolates were analyzed for intracellular localization using transmission electron microscopy (TEM), structured illumination microscopy (SIM), serial block-face scanning electron microscopy (SBF-SEM), and imaging flow cytometry. Transcriptional reprogramming of infected U937 monocytes was examined by mRNA sequencing. Infection outcomes were characterized and compared to A549 and SaOS-2 cell lines employing Luminex cytokine assays, flow cytometry and Western blot analysis to characterize host cell death mechanisms in both wild-type and TNF-R1 deficient backgrounds.

**Results:** All *S. aureus* isolates localized to endolysosomal and cytosolic compartments but also peri⍰ and putatively intranuclearly, revealing an unexpected intracellular niche. In U937 monocytes, infection induced a conserved stress signature alongside isolate⍰specific transcriptional programs divergently affecting inflammation, metabolism, and cell fate, which was markedly attenuated in response to the chronic⍰infection isolate EDCC 5464. Cell death outcomes were likewise isolate⍰dependent, involving intrinsic and extrinsic apoptosis, mitochondrial depolarization, and caspase-1 activation at distinct temporal dynamics. TNF⍰R1 loss initially delayed but exacerbated late, isolate-independent cytotoxicity, identifying TNF⍰R1 as a key regulator of U937 infection outcome. SaOS⍰2 and A549 cell death was far less affected by isolate or TNF-R1 deficiency.

**Conclusions:** These results highlight the multilayered determinants governing intracellular *S. aureus* survival, non-canonical intracellular localization, and host cell susceptibility. The TNF/TNF-R1 axis is identified to critically determine regulated host defense during early infection stages in a tissue-specific manner. Together with distinct isolate-driven gene expression profiles, infection risks under TNF-targeted therapies and the contribution of *S. aureus* heterogeneity should be considered in the design of future host-directed treatment strategies.

**Plain English summary:** The bacterium *Staphylococcus aureus* (*S. aureus*) often lives harmlessly in humans but can cause severe or recurrent infections when the skin barrier is broken or the immune system is weakened. A major reason for its persistence is its ability to hide inside human cells, where it is shielded from immune attacks and antibiotics. To effectively target such bacteria, it is crucial to understand that infections vary depending on both the bacterial strain and the infected cell type. Many reasons behind these differences are still puzzling. We explored how different types of *S. aureus* (collected from different disease types) change how human cells respond to infection. We focused on how the different strains influence the way immune cells adjust their gene activity during infection, and how a receptor called TNF-R1 is involved in managing cell death responses.

Bacteria were found not only in compartments meant to destroy them but also near and even inside the cell nucleus, an unexpected location. All strains triggered a similar stress response but also distinct patterns influencing inflammation, metabolism, and cell survival. A strain linked to chronic infection caused weaker responses, suggesting greater stealth. Cells lacking TNF-R1 initially survived longer but later showed greater damage, indicating this receptor’s role in infection control. In lung and bone cells, these effects were less pronounced.

Concludingly, *S. aureus* occupies unexpected niches inside human cells and uses varying survival strategies. TNF-R1 is a key regulator of host infection responses in the analyzed immune cells, highlighting that both bacterial diversity and host factors must be considered when developing targeted treatments.

**Graphical Abstract:** Peri- and intranuclear localization early after *S. aureus* uptake across host cell lines, with isolate-specific modulation of host fates and a critical role for TNF-R1 to mediate regulated death responses of U937 cells.
At 2 hpi, intracellular *S. aureus* not only localizes in (LAMP-1 decorated) membrane-enclosed compartments or directly in the cytosol, but within invaginations of the nuclear surface and intranuclearly with or without being surrounded by a vesicular membrane in U937_wt_, SaOS-2_wt_, and A549_wt_ cells. At 4 hpi, *S. aureus* triggers differential gene expression in (**A**) U937_wt_ cells to an isolate-specific extent, with both unique and shared transcriptomic signatures across the four isolates, that is muted for the chronic infection isolate EDCC 5464. Apoptotic cell death is induced to an isolate-dependent extent involving extrinsic initiator caspase-8, intrinsic initiator caspase-9 (EDCC 5055 only), and variable effector caspase-3/-7 activity in the earlier stages of infection (6 hpi), which then barely increases (24 hpi) in U937_wt_ cells. *S. aureus*-induced cell death and caspase activation is abolished in (**B**) U937_ΔTNF-R1_ at 6 hpi, but is significantly reinforced at 24 hpi with diminished isolate-specificity. Correspondingly, mitochondrial trans-membrane potential (ΔΨm) is disrupted for all isolates upon TNF-R1 knockout, as well as caspase-1 activity, suggesting pyroptotic pathway activation at later stages of infection. (**C**) SaOS-2 _wt_ cells show moderate caspase-3/-7 and -1 activation, while infection induces detachment of (**D**) A549_wt_ cells with minimal caspase activation. Infection induces an isolate- and cell line-dependent cytokine release. Coloured arrows indicate the mean proportion of effector-positive cells (↑ ∼20-40%, ↑ ↑ 40-60%, ↑ ↑ ↑ >60%) representing each *S. aureus* isolate. Grayed signaling arrows indicate the hypothesis by which TNF-R1 activation and internalization is required to kill lysosomal *S. aureus* via activation of anti-microbial enzymes and downstream regulated death pathway activation. Created with BioRender.com.

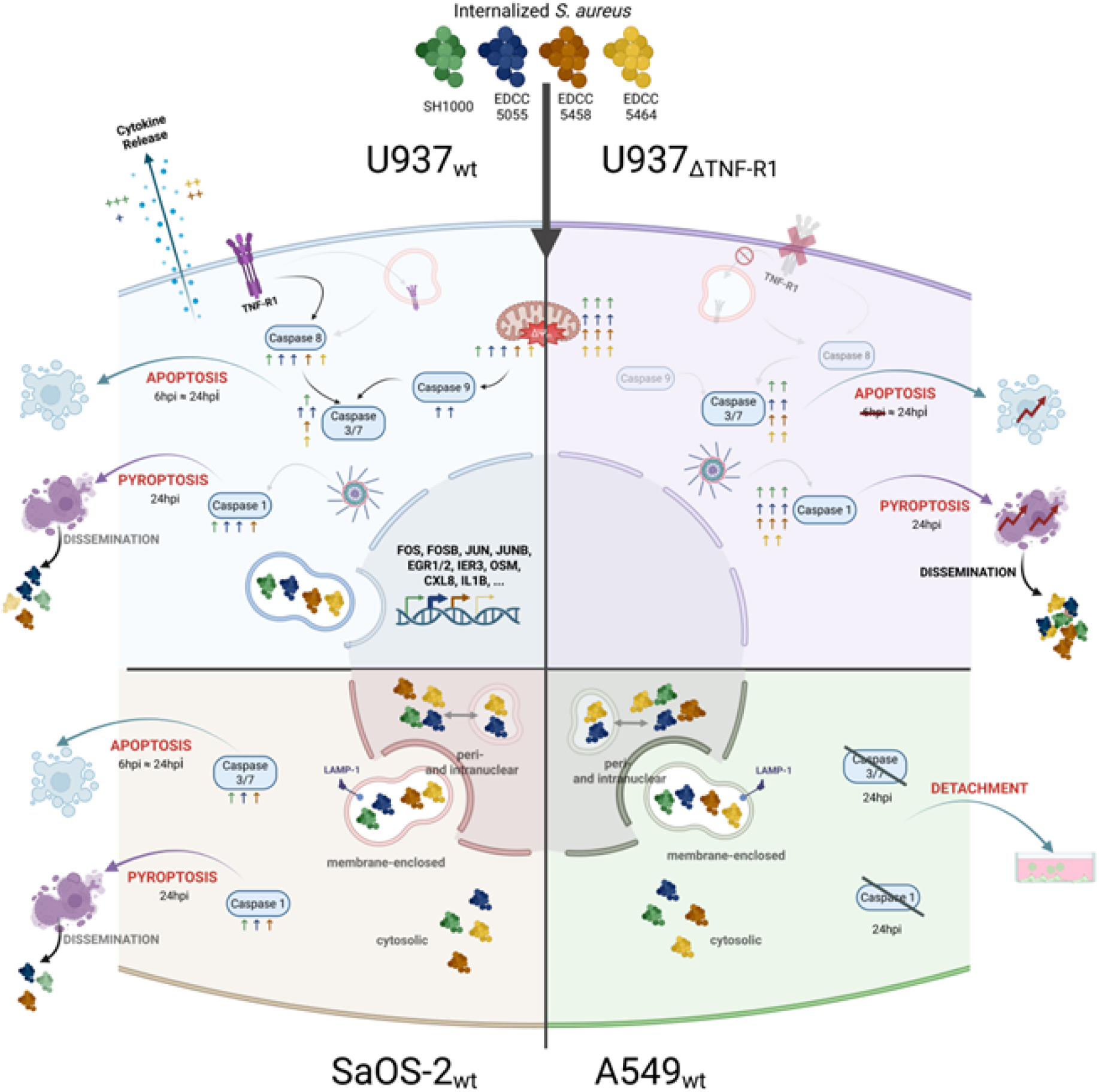

## Background

*Staphylococcus aureus* (*S. aureus*) asymptomatically colonizes up to one third of healthy individuals yet remains a major cause of community⍰ and hospital⍰acquired infections, ranging from superficial disease to severe pneumonia, endocarditis, osteomyelitis, and sepsis ^1,2^. Increasing evidence shows that a majority of *S. aureus* strains are proficient intracellular pathogens capable of invading, surviving, and persisting within diverse professional and non⍰professional phagocytic cell types. These include phagocytosis by professional phagocytes such as neutrophils, monocytes and macrophages, as well as internalization by non-professional phagocytic epithelial and endothelial cells, keratinocytes, and osteoblasts ^3-11^. This intracellular versatility is supported by high phenotypic plasticity and tightly regulated virulence programs, including Agr, SaeRS, SarA, or SigB ^12,13^, enabling adaptation to distinct tissue niches, evasion of immune surveillance, and persistence despite or even promoted by antibiotic treatment with limited intracellular activity. Consequently, intracellular survival critically contributes to chronic and therapy-refractory infections ^14,15^.

Infection outcomes, however, are shaped not only by strain⍰specific virulence patterns but also by the host cell type, resulting in highly specialized host-pathogen interactions. Professional phagocytes act as a first line of antimicrobial defence, but may act as “Trojan horses” during *S. aureus* systemic dissemination when bacterial clearance fails ^4,16-19^. In contrast, non-professional phagocytic cells may internalize bacteria less efficiently yet provide distinct intracellular niches that determine host and bacterial fate.

Once internalized, *S. aureus* disrupts phago-lysosomal trafficking and maturation, that can be followed by escape into the cytosol, and modulates cellular stress and inflammatory pathways. Manipulation of host cell death signaling is particularly decisive for whether intracellular bacteria are contained, eliminated, or are ultimately released in a manner that promotes inflammation and dissemination ^20-22^.

Central to this balance is the activation of the tumor necrosis factor-α (TNF) / TNF receptor 1 (TNF-R1) system, which coordinates inflammation, antimicrobial defence, and regulated cell death. However, excess TNF-signaling can potentiate cell death to an extent of cellular demise and severe tissue damage ^23^. Formation of signaling *complex I* on plasma membrane resident receptors triggers downstream activation of NF-κB and MAP kinases (MAPK) to promote cell survival, proliferation, and activation of cytokine signaling on the one hand, while re-organization of *complex I* to *complex IIa/b* on the other hand results in the activation of apoptotic cell death cascades ^23,24^. In case of absent or inactive caspase-8, signaling shifts towards necroptosis ^25^. In some cell types, full activation of TNF-mediated apoptosis requires endocytosis of complex-II bearing TNF-R1 ^26^. Indeed, several bacteria have evolved mechanisms to subvert this axis by inhibiting TNF-R1 internalization or endo-lysosomal fusion processes, thereby preventing lysosomal degradation and apoptosis to maintain their replicative niche ^27-30^.

Clinically, blockade of TNF signaling may be beneficial to attenuate excessive inflammation and severe tissue damage by overshooting immune responses during infection. However, anti-TNF treatment in auto-immune diseases increases susceptibility to and recurrence of infections by opportunistic and intracellular pathogens, including *S. aureus*, due to impaired TNF-dependent immune defences necessary for adequate bacterial clearance ^31-34^. Thus, balancing anti-inflammatory benefit with infection risk requires a precise understanding of TNF-R1-mediated control of intracellular *S. aureus* survival, host cell death modalities, and tissue-specific outcomes.

Despite extensive work on intracellular *S. aureus*, comparisons across strains and host cell types remain limited, complicating generalizable conclusions beyond the specific model used in each study ^6,9^, To address this, we systematically investigated host responses of the human monocytic cell line U937 infected with the well-characterized *S. aureus* isolate SH1000 alongside three clinical strains of different, sepsis- and acute or chronic implant-associated bone infection backgrounds (EDCC 5055, EDCC 5458, and EDCC 5464) ^35-38^ and compared infection outcomes with those of non-professional phagocytic lung-epithelial and osteoblast-like A549 and SaOS-2 cell lines.

All isolates were internalized into membrane-enclosed compartments, along with cytosolic and, notably, peri- and intranuclear localization. Despite a conserved stress response across *S. aureus* isolates, each induced specific host transcriptional alterations in U937 monocytes, consistent with heterogeneous cytokine release and cytotoxicity. Host cell death involved isolate-dependent activation of apoptotic and non-apoptotic caspases at different stages of infection in U937 and SaOS-2 cells, while A549 cells showed minimal caspase activation despite isolate-specific detachment. TNF-R1 deletion in U937 cells abolished early regulated cell death but exacerbated late cytotoxicity across isolates, suggesting a critical role for TNF-R1 in the finely controlled balance of regulated host defence and deleterious cytotoxicity in U937 monocytes, but less in A549 and SaOS-2 cells.

Together, these findings reveal pronounced isolate- and tissue-specific responses in intracellular *S. aureus* infection and identify TNF-R1 as a critical determinant of monocyte infection outcomes, emphasizing the need for tissue-directed therapeutic strategies and careful consideration of TNF-modulating treatments in the context of intracellular bacterial infections.

## Methods

### Human cell lines, bacterial strains and culture conditions

U937_wt_ /U937Δ_TNF-R1_ ^103^, A549_wt_ / A549Δ_TNF-R1_ , and SaOS-2_wt_ / SaOS-2Δ_TNF-R1_ cells were cultivated in RPMI (Gibco), DMEM (Gibco), and McCoy’s (Capricorn Scientific), respectively, supplemented with 10% FBS (Gibco) and 1% sodium-pyruvate (Gibco) without antibiotics at 37°C, 5% CO_2_.

*Staphylococcus aureus* (*S. aureus*) strains EDCC 5055, EDCC 5458, EDCC 5464, and SH1000 ^35-38^ (kindly provided by GK. Mannala and S. Muehlen) were grown in Luria-Bertani broth (LB) aerobically at 37°C and 180 rpm overnight. The bacterial overnight cultures were adjusted to the stationary growth phase at an optical density at 600 nm (OD_600_) of 1.0-1.5 and incubated for at least 30 min at 37°C and 180 rpm. To calculate the multiplicity of infection (MOI), the bacterial number was determined with an OD_600_ of 1 corresponding to 8 × 10^8^ colony-forming units (CFU) per ml.

#### TNF-R1 knock-down

For CRISPR-mediated knockdown, we used commercial Cas9 (IDT) loaded with crRNA:tracrRNA duplexes as recommended by the manufacturer’s protocol (Alt-R CRISPR-Cas9, IDT).

Two TNF-R1 gene targeting, predesigned crRNAs (Hs.Cas9.TNFRSF1A.1.AA: TAATGTATCGCTACCAACGG; Hs.Cas9.TNFRSF1A.1.AC: AGAGGTGCACGGTCCCATTG) were used to enhance knockout performance, as recommended ^104^. Complexes were delivered by nucleofection (Amaxa), followed by Western blot screening.

### Intracellular infection assays

Unless stated otherwise, A549 and SaOS-2 cells were seeded at a density of 5×10^5^ cells/mL overnight, and U937 cells were adjusted to 1×10^6^ cells/mL prior to infection. Cells were infected with live or heat-killed (10 min, 95°C) *S. aureus* at MOI 30 or MOI 100, followed by 1 h of co-culture at 37°C, 5% CO_2_ to allow for synchronized intracellular uptake. To remove extracellular bacteria, cells were washed with PBS and resuspended in fresh medium containing gentamicin (G1397; Sigma-Aldrich) at 37.5 µg/mL to focus investigation of infection-induced effects by intracellular *S. aureus* only (Gentamicin protection assay) ^36^.

### Cell death analysis

For each assay, untreated cells served as negative controls. For positive apoptosis controls, cells were primed with cycloheximide (2.5 µM; API-03, VWR) 30 min prior to stimulation with TNF (100 ng/mL; provided by D. Männel, UKR) or stimulated with nigericin (20 µM; Cay11437-5, Biomol) for caspase-1 activation indicating pyroptotic pathway activation. At the indicated time points, staining was performed following the manufacturer’s instructions, unless stated otherwise. Caspase-3/7 and multiple-caspase assays were measured using Guava MUSE (Cytek). Annexin-V, caspase-1, -8, and -9 assays using a Guava easyCyte (Cytek) and guavaSoft (4.5.25, Cytek) for data analysis.

*Annexin V/7-AAD staining:* in brief, 250 µl of carefully resuspended U937 cells were sedimented (500 x g, 5 min), resuspended in 20 µl staining mix (18 µl PBS, 1 µl FITC-Annexin V, 1 µl 7-amino-actinomycin D (7-AAD); #640922, BioLegend), and incubated for 20 min in the dark. 100 µl of binding buffer were added to the samples prior to measurement.

#### Caspase Assays

For Muse® Caspase-3/7 (#MCH100108, Cytek) and Multi-Caspase assays (#MCH100109, Cytek), 50 or 100 µl of carefully resuspended cells were sedimented (following detachment in case of adherent cells), washed with PBS (5 min, 500 x g), and the pellet was stained following the manufacturer’s recommendations. In brief, each pellet was resuspended in 50 µl 1x assay buffer, 5 µl caspase-working solution added (substrate diluted 1:8 in PBS for Caspase-3/-7 or 1:160 for Multi-Caspase), and incubated for 30 min under cell culture conditions, followed by incubation with 148 µl assay buffer and 2 µl 7-AAD in the dark for 5 min at room temperature.

To assess caspase-1 positive cells, the FAM-YVAD-FMK substrate (#13483, Biomol) was diluted 1:250 in 250 µL cell suspension with 5 µL 7-AAD (#420404, BioLegend), followed by 1h incubation under cell culture conditions. Cell suspensions were washed once in PBS (5 min, 500 x g), resuspended in 100 µl PBS, and immediately analyzed. For Caspase-8 (FAM-LETD-FMK (656), #ICT099, BioRad) and Caspase-9 (FAM-LEHD-FMK (677), #ICT912, BioRad), 10 µl of caspase working solution (1:150 in PBS) were added to 290 µl of the cell suspension. After 60 min of incubation under cell culture conditions, cells were washed twice using 1x Apoptosis Wash Buffer (AWB) and resuspended in 300 µl 1 x AWB. Thereafter, samples were incubated with 1,5 µl (0.5% v/v) of propidium iodide (PI) for 5 min in the dark at room temperature. All samples were kept on ice and analyzed immediately after staining.

As an essential indicator of cellular health, stress, and death, changes in mitochondrial trans-membrane potential (ΔΨ_m_) along with cell membrane permeabilization were monitored using the Muse® MitoPotential assay (#MCH100110, Cytek) following the manufacturer’s instructions. In brief, the MitoPotential working solution was prepared at 1:1,000 in 1x Assay Buffer, 95 µl were added to 100 µl of the cell suspension and incubated for 20 min under cell culture conditions. Following incubation with 5 µl of 7-AAD in the dark for 5 min at room temperature, samples were analyzed immediately.

#### SDS-PAGE and Western Blot

After the indicated time points, cells were sedimented and washed with PBS (5 min, 500 x g, 4°C). Cell pellets were lysed in 30 µl modified RIPA buffer (50 mM TRIS-HCl [pH 7.5], 150 mM NaCl, 1% NP-40, 1% Triton X-100, 1 mM EDTA, 0.25% Na-deoxycholate), containing a protease inhibitor cocktail (4693124001, Sigma-Aldrich), followed by 20 sec sonication (cooled to 4°C) and 5 min sedimentation at 1500 x g at 4°C. Supernatant was used for BCA protein quantification (Pierce, #23225). Samples were prepared immediately before SDS-PAGE using 12.5% or 10% PAA gels. Proteins were blotted onto a PVDF membrane (200T.1, Carl-Roth). The membranes were blocked with 5% skimmed milk in TBS-T and incubated overnight with the primary antibodies, diluted 1:1,000 in 5% skimmed milk. The peroxidase-conjugated secondary antibodies were incubated for 1 h, diluted 1:10,000 in TBS-T. Blots were imaged using ECL (RPN2232, Cytiva) and the ImageQuant800 device (Cytiva). Primary antibodies: PARP-1 (cs #9542S), cleaved caspase-3 (#9661S, Cell Signaling), caspase-3 (#9668S, Cell Signaling), Actin-HRP (HRP-60008, Proteintech). Secondary antibody: anti-rabbit-HRP (111–035–144, Jackson ImmunoResearch).

#### xCELLigence Real Time Cell Analysis (RTCA)

To analyze both the extent and kinetics of cell death of adherent cells, xCELLigence Real-time cell analysis (RTCA, OMNI Life Science) assay was performed following the manufacturer’s instructions. In brief, A549 or SaOS-2 cells were seeded overnight at 8000 cells per well in 100 µL medium after a blank measurement using 50µL of the corresponding cell culture medium. The attached cells were then infected in duplicates with the corresponding *S. aureus* isolates at MOI 30 and co-cultured for 1h. Along with a duplicate of entirely untreated cells, both uninfected and infected cells were then washed thrice in and resupplied with fresh cell culture medium containing 37.5 µg/ml gentamicin (G1397; Sigma-Aldrich). The detachment of cells based on cellular impedance (indicating cell death) was monitored every 15 minutes for at least 48 hours post infection (hpi).

### Luminex Multiplex ELISA®

To monitor the release of selected cytokines, 1×10^6^ U937 cells or 5×10^5^ A549 cells were infected with *S. aureus* at MOI 30. At 4 hpi, cells were centrifuged at 14,000 xg for 10 minutes. A 48-plex Luminex Multiplex ELISA (Bio-Rad Laboratories, #12007283) was performed using the supernatants according to the manufacturer’s protocol ^105^, using the Bio-Plex 200 system with HTF (#171000205, Bio-Rad Laboratories). Thereupon, the assay was repeated using a customized 8-plex Luminex Multiplex ELISA (IL-6, IL-8 (CXCL8), MCP-1 (CCL2), MIF, MIP-1α (CCL3), RANTES (CCL5), SCGFβ, TNF-α) (#PPX-08, Thermo Fisher) in all three cell lines.

### Labelling of bacteria

For Lipobiotin/Streptavidin-labelling we followed ^106^. The bacterial suspension was centrifuged (10 min, 10,000 x g) and the sediment was resuspended in 300 µl PBS. 5 µg/ml of lipobiotin (L2070; EMC microcollections) was added, together 10 µl of Streptavidin-magnetic nanobeads (130-048-102; Miltenyi Biotec). After incubation at 37°C for 1 h, bacteria were washed once in PBS (10 min, 10,000 x g), then resuspended in 500 µl PBS. The number of labelled bacteria was determined photometrically at OD_600_ and used for infection.

### Transmission Electron Microscopy (TEM)

Adherent cells were seeded on Corning™ BioCoat™ fibronectin-coated coverslips (Fisher Scientific, #08-774-386) one day and U937 suspensions cells directly prior to infection in crowded conditions in a 1.5 ml tube. Cells were infected with either two (A549, SaOS-2, n=2) or four (U937, n=1) of the *S. aureus* isolates at MOI 100, respectively, which were labelled with lipobiotin and streptavidin-conjugated magnetic nanobeads (130-048-102; Miltenyi Biotec), which will be used for targeted organelle purification in future investigations.

Following a washing step and gentamicin treatment at 1 hpi, cells were fixated at 2 hpi in 2% glutaraldehyde (Serva Electrophoresis) buffered with 100 mM sodium cacodylate, pH 7.4 for 1 h at room temperature. Afterwards the suspension cells were coated in 4% low melting agarose (840302; Biozym), dissolved at 37°C in 100 mM sodium cacodylate and then cooled for 35 min on ice. After several washing steps with 100 mM sodium cacodylate, all cell lines were further fixated using 0,5% osmium tetroxide (Science Services) and 0.5% uranyl acetate was added for optimal contrast. Both treatments were performed for 30 min at 4°C. The cells were dehydrated in an ascending series of ethanol and finally acetone on ice. For equilibration, the specimens were treated overnight at 4°C in a 1:1 solution of Epon (Fluka) and acetone, followed by embedding with pure Epon for 2 h at 30°C and polymerization for 2 d at 60°C. For adherent cell samples, cover slips were removed without damaging the embedded cells by alternately exposing them to liquid nitrogen and heated water (60 °C). Ultrathin sections were cut with a diamond knife (Diatome) on an ultramicrotome EM UC7 (Leica) and collected on a hexagonal 200 mesh copper grid (Plano). The electron micrographs were taken at 80 kV on a Zeiss 902 or at 60 kV on a Zeiss EM10a transmission electron microscope (Zeiss). Several cells of different ultrathin sections were captured at different magnitudes and TEM pictures were optimized for brightness and contrast using ImageJ ^107^.

### Serial block-face scanning electron microscopy (SBF-SEM)

Cultured cells grown on 8-well chambered coverslips (IBIDI) were fixed in 0.1 M cacodylate buffer (pH 7.4, Science Services, #E11652) containing 2.5% glutaraldehyde (Science Services, #E16216). Samples were subsequently processed following a modified NCMIR rOTO-post-fixation protocol ^108^, which enhances contrast and confers resistance to electron beam damage during serial imaging of EPON resin-embedded specimens.

### Imaging Flow Cytometry

Bacteria were labeled with 5 µM CFSE/PBS for 1 h, shaking at 37°C, followed by three washing steps (10,000 × g, 10 min) in PBS. 1×10^6^ U937 cells were infected with CFSE-labeled bacteria for 2 h at MOI 100. At 1 hpi, gentamicin was added to kill extracellular bacteria. 5 min before the end of the infection period, Cell Mask® deep red membrane stain (C10046; Thermo Fisher) was added, diluted 1:25,000 in PBS, and incubated for 5 min, followed by washing twice in cold PBS. Cells were then fixed in 4% PFA/PBS for 15 min on ice, followed by washing in 1 ml cold PBS. Cells were stored in the dark and cooled until imaging using ImageStream MKII (Cytek). 10,000 images per condition were acquired at 60x using the Inspire software, capturing channels Ch01 (Brightfield), Ch02 (CFSE), and Ch11 (CellMask deep red). Images were analyzed using Ideas software (V6.2.65): Cells were gated: in-focus/single cells/double positive. Then, composites of Ch02/Ch11 were generated.

### SIM imaging

SIM was performed on a Zeiss Elyra 7 equipped with a C-Apochromat 63x/1.4 NA oil-immersion objective and following laser for excitation: high-reflectivity (HR) Diode 405 nm (50 mW), HR Diode 488 nm (500 mW), HR diode-pumped solid state 561 nm (500 mW), and HR Diode 642 nm (500 mW). Emission light was passed through these dichroic filter sets: BP420-480+LP655 for the DAPI and Alexa Fluor 647 channel and BP495-525+LP655 for the Alexa Fluor 488 channel. Emission light was further split by a SBS LP 560 beam splitter and detected on two PCO Edge 4.2M sCMOS camera sensors. For 3D-SIM imaging appropriate grid-settings, laser lines and filter combinations were chosen following the suggested settings by the ZEN black microscope control software. Z-stacks were acquired using the Z-step interval optimized for “3D leap mode” as defined by ZEN, with 13 phase shifts recorded per z-plane. 3D-SIM reconstruction was performed in ZEN black using the standard SIM processing preset for fixed samples. Chromatic aberration was corrected by imaging a multicolor bead sample (T14792, Thermo Fisher) using identical imaging and processing settings. A channel alignment matrix was then generated in ZEN black and applied during the SIM processing of the sample images.

### Post-processing of SIM image

SIM images were processed in Imaris 11.0.0 (Oxford instruments) using the spot detection and surface generation function. Fist, nuclei were first segmented using the Surface Generation tool with the integrated machine-learning–based training workflow. Nuclear surfaces were subsequently filtered by voxel count to remove segmentation artifacts and nuclei not fully contained within the field of view. Bacterial DNA stained by DAPI was detected using the Spot Detection function with an estimated diameter of 0.42 µm and background subtraction enabled. Detected spots were filtered by Imaris “Quality” parameter (> 1500) and further curated with a machine-learning assisted step to exclude false positive detections from high heterochromatin densities within the cellular nucleus. LAMP-1 signal was segmented using Surface function and filtered to only include surfaces with >2000 voxels. The shortest distances from detected bacteria to the nucleus and Lamp1 surface were then extracted to quantify bacteria localization and LAMP-1–spot proximity.

### Quantification of intracellular bacteria – colony-forming units

Cells were infected as described previously. At the indicated time points, cells were washed with PBS (500 x g, 5 min for U937 cells) to remove bacteria that had been released from the host cells. Thereafter, cells were lysed using 100 µl or 500 µl of 1% Triton X-100 (in ddH_2_O) for suspension cells and adherent cells, respectively, and incubated at 37°C for 10 min. Following thorough vortexing and resuspension, the lysate was serially diluted, 10 µL of the dilution were plated on LB-agar plates and incubated overnight. CFU were calculated as follows:

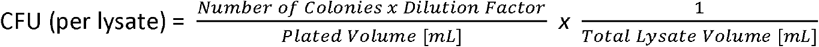

### RNA sequencing and data analysis

To screen for differential gene expression upon infection, U937 cells were infected with four different isolates of *S. aureus* or, for comparison, vancomycin-resistant *E. faecium* for 4 or 6 h, respectively, or left untreated. RNA was isolated using RNAplus kit (Qiagen). Library preparation and RNAseq were carried out as described in the Illumina “Stranded mRNA Prep Ligation” Reference Guide, the Illumina NextSeq 2000 Sequencing System Guide (Illumina), and the KAPA Library Quantification Kit - Illumina/ABI Prism (Roche Sequencing Solutions, Inc., Pleasanton, CA, USA). 200 ng of total RNA was used for purifying the poly-A-containing mRNA molecules using oligo(dT) magnetic beads. Following purification, the mRNA was fragmented to an average insert size of 200-400 bases using divalent cations under elevated temperature (94°C for 8 min). Next, the cleaved RNA fragments were reverse transcribed into first-strand complementary DNA (cDNA) using reverse transcriptase and random hexamer primers. Thereby, Actinomycin D was added to allow RNA-dependent synthesis and to improve strand specificity by preventing spurious DNA-dependent synthesis. Blunt-ended second-strand cDNA was synthesized using DNA Polymerase I, RNase H, and dUTP nucleotides. The incorporation of dUTP, in place of dTTP, quenches the second strand during the later PCR amplification, as the polymerase does not incorporate past this nucleotide. The resulting cDNA fragments were adenylated at the 3’ ends, and the pre-index anchors were ligated. Finally, DNA libraries were created using a 13 cycles PCR to selectively amplify the anchor-ligated DNA fragments and to add the unique dual indexing (i7 and I5) adapters. The libraries were bead-purified twice and quantified using the KAPA Library Quantification Kit. Equimolar amounts of each library were sequenced on an Illumina NextSeq 2000 instrument controlled by the NextSeq 2000 Control Software (NCS) v1.7.0.45751, using one 50 cycles P3 Flow Cell with the dual index, single-read (SR) run parameters and the dark cycle recipe. Image analysis and base calling were done by the Real Time Analysis Software (RTA) v4.12.2. The resulting .cbcl files were converted into fastq files using bcl2fastq v2.20 software. Library preparation and mRNAseq were performed at the Genomics Core Facility “KFB - Center of Excellence for Fluorescent Bioanalytics” (University of Regensburg, Regensburg, Germany; www.kfb-regensburg.de).

Processing of raw data was performed at the Core Unit Systems Medicine, University of Würzburg (Würzburg, Germany; https://www.med.uni-wuerzburg.de/cu/sysmed/). To assure high sequence quality, Illumina reads were quality- and adapter-trimmed via Cutadapt ^109^ v2.5 using a cutoff Phred score of 20 in NextSeq mode, and reads without any remaining bases were discarded (parameters: -- nextseq-trim=20 -m 1 -a ACTGTCTCTTATACACATCT). Processed reads were subsequently mapped to the human genome (NCBI RefSeq assembly GCF_000001405.40 / GRCh38.p14, primary assembly and mitochondrion) using STAR ^110^ v2.7.2b with default parameters but including transcript annotations from RefSeq annotation version RS_2023_03 for GRCh38.p14. This annotation was also used to generate read counts on exon level summarized for each gene via featureCounts v1.6.4 from the Subread package ^111^. Multi-mapping and multi-overlapping reads were counted strand-specific and reversely stranded with a fractional count for each alignment and overlapping feature (parameters: - s 2 -t exon -M -O --fraction). The count output was utilized to identify differentially expressed genes using DESeq2 ^112^ v1.24.0. Read counts were normalized by DESeq2 and fold-change shrinkage was conducted by setting the parameter betaPrior to TRUE. Separate DESeq2 analyses were conducted for comparisons within *S. aureus* and *E. faecium* infection data, both including the uninfected control.

Principal component analysis (PCA) was performed by applying variance-stabilizing transformation (vst) to normalized read counts via DESeq2. The 1000 most variable genes (after removal of zero/NA-variance features) across all samples were selected and scaled prior to PCA computation using R (prcomp, stats package; R version 4.5.1, RStudio 2025.5.1.513). Samples were grouped by condition (9 levels: SH1000, EDCC 5055, EDCC 5458, EDCC 5464, untreated, and four *E. faecium* strains VRE 23×1608, VRE 23×1609, VRE 7872, and VRE 8170) and PCA scores were plotted using ggplot2 (version 3.5.2).

Gene lists used for all downstream visualization and functional analyses were defined as protein-coding, significantly regulated (adjusted p-value after Benjamini-Hochberg correction (padj) < 0.05), and at least two-fold up- or downregulated (|log2FC| ≥ 1) relative to uninfected control cells or between cells infected with different *S. aureus* isolates, as applicable. Results shown here focus on *S. aureus*-infected cells. Full STRING networks (https://string-db.org/) were generated at medium confidence (0.400), with edges indicating confidence and disconnected nodes removed. Heat maps of selected genes were created using GraphPad Prism 10.6.1.

Functional enrichment of biological pathways that are present in these gene lists more than expected by chance was performed with g:Profiler ^113^ (version e113_eg59_p19_f6a03c19). Only annotated genes were submitted as ordered queries using the g:SCS multiple-testing correction and a significance threshold of 0.05, allowing the analysis to account for the ranked structure of the input lists without imposing additional arbitrary cutoffs. Gene Ontology Biological Process (GO:BP) terms were used for all downstream analyses. For ordered queries, the adjusted values reported by g:Profiler do not represent classical *P*-values but are to be interpreted as enrichment scores (adj_score) that reflect the concentration of term-associated genes at the top of the ranked list rather than random distribution ^114^.

To identify non-redundant groups of related GO:BP terms, enriched up- and downregulated pathways were imported separately for each condition into R (R version 4.5.1, RStudio 2025.5.1.513) and processed with *tidyverse* (version 2.0.0) and *openxlsx* (version 4.2.8). For every term, the associated gene set was represented as a vector of gene symbols derived from the g:Profiler “intersections” column. Gene-set similarity was quantified as the Jaccard coefficient (intersection/union of member genes), as commonly applied in gene-set clustering tools such as EnrichmentMap and GOMCL ^115-117^. Hierarchical clustering (*hclust*, stat package) was performed based on a Jaccard distance matrix (1 − Jaccard coefficient) between pathways using average linkage. To obtain interpretable clusters with comparable granularity across isolates, dendrograms were cut at height h=0.5 (*cutree*), corresponding to a minimum within-cluster Jaccard similarity of 0.5, consistent with conservative Jaccard thresholds recommended for pathway network construction. Small or single-pathway clusters were retained, as the goal was to preserve biologically specific terms rather than impose arbitrary minimum cluster sizes. Each cluster was named manually according to the driver term. The minimum adjusted enrichment score (min_adj_score) among its member pathways and the total number of genes of each cluster were extracted and visualized as bubble plots in R using ggplot2 (*version 3*.*5*.*2*), with bubble size encoding gene count per cluster and bubble color representing −log10 of the cluster-level min_adj_score, similar to standard enrichment bubble plots. R scripts for pathway clustering and visualization are provided as **Supplementary Data S1**.

### Sequencing, assembly, and annotation of S. aureus genomes

Four isolates were sequenced using both short-read Illumina and long-read Oxford Nanopore technologies. Illumina sequencing was performed on a NextSeq platform using Illumina DNA Prep chemistry (mid-output, 300 cycles, paired-end 2 × 149 bp). Long-read sequencing was conducted on a GridION device using R10.4.1 flow cells and the Native Barcoding Kit 96 v14 (SQK-NBD114.96). Hybrid genome assemblies were generated from the corresponding FASTQ files using MiLongA version 1.0.3 ^118^. For assembly generation and quality evaluation, only the components relevant for Flye-based assembly were applied, namely Flye v2.9.4-b1799 for long-read assembly, minimap2 v2.28-r1209 for read mapping, and CheckM v1.2.2 for genome completeness and contamination assessment. The Flye assemblies were selected for all downstream analyses. The resulting genome assemblies were annotated using the NCBI Prokaryotic Genome Annotation Pipeline (PGAP, release 2023-10-03.build7061) ^119^ and are available in **Supplements SX-SX**/NCBI.

To compare genomes between the different isolates (**Supplementary File S2**), assembled data were annotated using Prokka (1.14.6; ^120^) and comparison tables were generated using ROARY (3.13.0; ^121^).

### Proteomic profiling of S. aureus at the steady-state

#### Filter aided sample preparation (FASP)

For the proteome analysis, samples were prepared using a modified filter-aided sample preparation (FASP) as detailed before ^122,123^. In brief, *S. aureus* isolates were lysed adding 700 µL of lysis buffer containing 7 M urea, 2 M thiourea, 5 mM dithiothreitol (DTT), 2% (w/v) CHAPS. Lysis was promoted by sonication using a Branson sonifier at output 5 for 1 min, followed by centrifugation at 10,000 x g for 10 min to pellet remaining bacterial cells.

Protein concentration of the supernatants was determined using the Pierce 660 nm protein assay (Thermo Fisher Scientific) following the manufacturer’s protocol. Proteins (corresponding to 20 µg) were transferred onto spin filter columns (Micron-30 Centrifugal Filters; Merck Millipore). Subsequently, samples were washed three times with a buffer containing 8 M urea. After reduction and alkylation by DTT and IAA, excess IAA was quenched with DTT and the membrane washed three times with 50 mM NH_4_HCO_3_. Afterwards, proteins were digested overnight at 37°C with trypsin (Trypsin Gold, Promega) using an enzyme-to-protein ratio of 1:50 (w/w). After digestion, peptides were recovered by centrifugation and two additional washes with 50 mM NH_4_HCO_3_. Combined flow-throughs were acidified with TFA to a final concentration of 1% (v/v) TFA and lyophilized. Purified peptides were reconstituted in 0.1% (v/v) FA for LC-MS analysis.

#### Liquid-chromatography mass spectrometry (LC-MS)

Proteome samples were analyzed using a nanoElute LC system (Bruker Corporation) coupled online to a timsTOF HT mass spectrometer (Bruker Corporation, Billerica, MA, USA). Peptides (corresponding to 200 ng) were separated at 400 nl/min using a reversed phase C18 column (Aurora ULTIMATE UHPLC emitter column, 25 cm x 75 µm 1.7 µm, IonOpticks), which was heated to 50°C. Peptides were loaded onto the column in direct injection mode at 600 bar. Mobile phase A was 0.1% FA (v/v) in water and mobile phase B 0.1% FA (v/v) in ACN. Peptides were separated running a linear gradient from 2% to 37% mobile phase B over 39 min. Afterwards, column was rinsed for 5 min at 95% B. Eluting peptides were analyzed in positive mode ESI-MS using parallel accumulation serial fragmentation enhanced data-independent acquisition (diaPASEF) ^124^. The dual TIMS was operated at a fixed duty cycle close to 100% using equal accumulation and ramp times of 100 ms each spanning a mobility range from 1/K_0_⍰=⍰0.6 Vs⍰cm^−2^ to 1.6 Vs⍰cm^−2^. We defined 36 × 25⍰Th isolation windows from *m/z* 300 to 1,165 resulting in 2-3 diaPASEF scans per TIMS cycle and an overall cycle time of 1.7 s. The collision energy was ramped linearly as a function of the mobility from 59 eV at 1/K_0_⍰=⍰1.3 Vs cm^−2^ to 20 eV at 1/K_0_⍰=⍰0.85 Vs cm^−2^. Samples were measured in triplicates.

#### Raw data processing

MS raw data were processed separately for each isolate using DIA-NN (version 2.2.0) ^125^ applying the default settings for library-free database search. Custom databases were compiled using isolate transcript libraries as described above as well as a list of common contaminants. Trypsin was set as the protease for in-silico library generation and peptide identification, allowing one missed cleavage. Carbamidomethylation was set as fixed modification and maximum number of variable modifications was set to zero. Peptide length ranged between 7-30 amino acids. The precursor m/z range was set to 300-1,800 and the product ion m/z range to 200-1,800. “QuantUMS (high precision)” mode was applied as quantification strategy with RT-dependent normalization. MS1 and MS2 mass accuracies were both set to 15 ppm and scan window size was automatically optimized using the build in algorithm of DIA-NN. Peptides precursor FDRs were controlled below 1%.

Proteins had to be identified by at least two peptides to be included in the final dataset.

#### Data availability

RNA-seq data are available at https://www.ncbi.nlm.nih.gov/geo/query/acc.cgi?acc=GSE328149.

The mass spectrometry proteomics data have been deposited to the ProteomeXchange Consortium (http://proteomecentral.proteomexchange.org) via the jPOST partner repository ^126^ with the dataset identifiers PXD073241 (ProteomeXchange) and JPST004317 (jPOST).

To review the data, go to: https://repository.jpostdb.org/preview/818370539696e370e1de6c with the access key: 2097

### Statistical analysis

Statistical analysis was performed using GraphPad Prism 10.6.1 software (GraphPad Software Inc.). Normality and Long-normality Tests were performed using the Shapiro-Wilk test. One- or Two-Way ANOVA with Tukey’s or Kruskal-Wallis correction for multiple comparisons were performed for testing of multiple groups, where appropriate. Comparisons were made for infected vs. uninfected control cells, between cells infected with different isolates or between cell types (wild-type versus knock-out or cells with versus without bc), as indicated respectively. A p value of <0.05 was considered statistically significant.

## Results

### Different *S. aureus* isolates are internalized by U937, A549, and SaOS-2 cells, showing not only lysosomal and cytosolic, but peri- and intranuclear localization early after uptake

Prior to internalization, we compared the *S. aureus* strains SH1000, EDCC 5055, EDCC 5458, and EDCC 5464 ^35-37^. Genome sequencing and steady⍰state proteomics (1h culture in cell culture medium) revealed a shared core virulence repertoire across isolates prior to host cell contact (**Table 1; Supplementary File S2**), including major adhesins (FnBPA, ClfA, ClfB), pore⍰forming toxins (α-hemolysin, LukGH), EsxA, staphopain A, or the regulators SrrA and SaeS. EDCC 5464 lacked detectable PSM⍰α/β/δ and several Spl proteases, consistent with reduced lytic activity and a persistent phenotype, while EDCC 5055 uniquely expressed MecA.

**Table 1.**
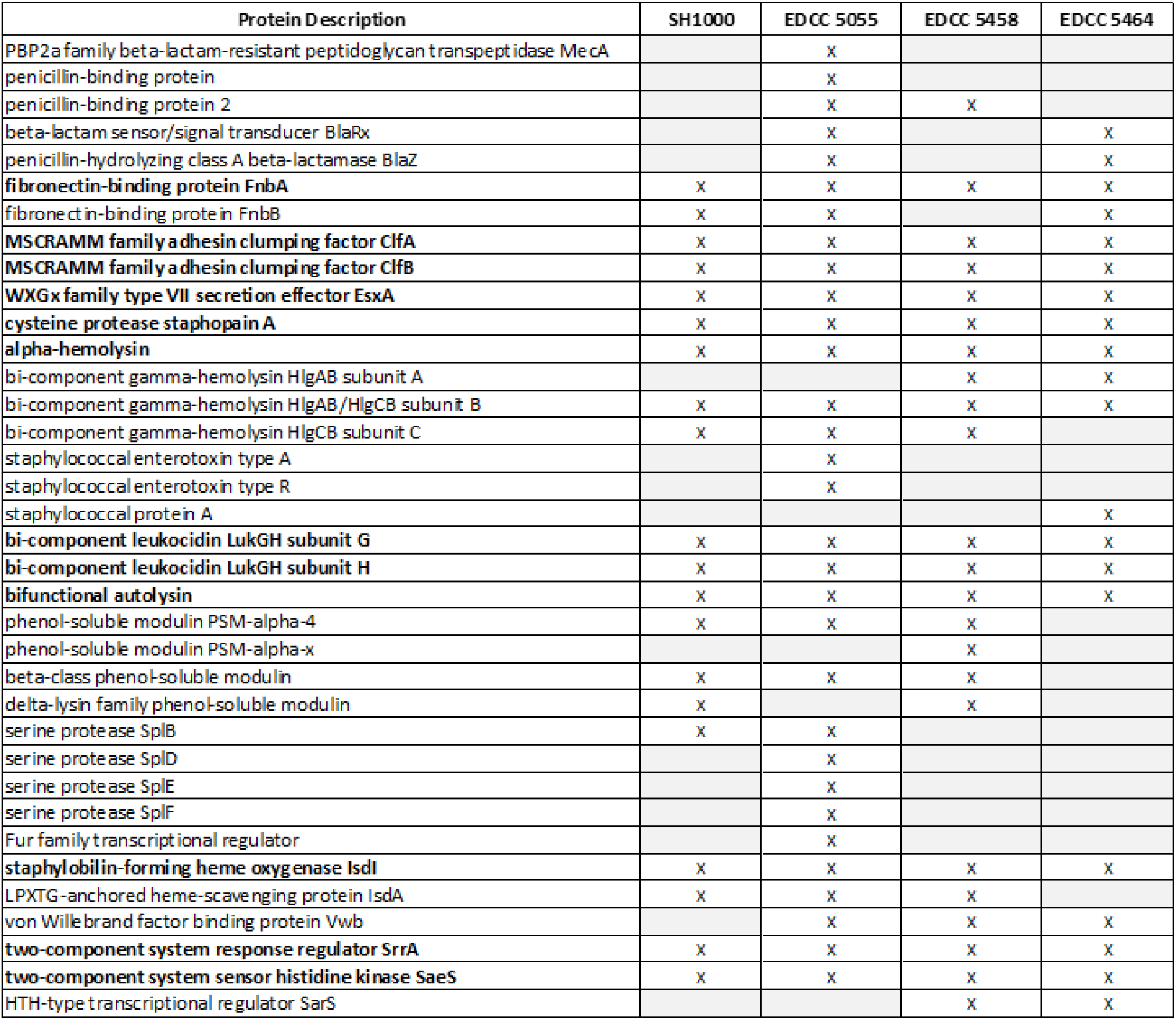
*S. aureus* steady-state expression of selected proteins prior to infection of host cells present with at least two peptides. Proteins expressed by all four isolates are highlighted (bold).

| Protein Description | SH1000 | EDCC 5055 | EDCC 5458 | EDCC 5464 |
| --- | --- | --- | --- | --- |
| PBP2a family beta-lactam-resistant peptidoglycan transpeptidase MecA |  | X |  |  |
| penicillin-binding protein |  | X |  |  |
| penicillin-binding protein 2 |  | X | X |  |
| beta-lactam sensor/signal transducer BlaRx |  | X |  | X |
| penicillin-hydrolyzing class A beta-lactamase BlaZ |  | X |  | X |
| fibronectin-binding protein FnbA | X | X | X | X |
| fibronectin-binding protein FnbB | X | X |  | X |
| MSCRAMM family adhesin clumping factor ClfA | X | X | X | X |
| MSCRAMM family adhesin clumping factor ClfB | X | X | X | X |
| WXGx family type VII secretion effector EsxA | X | X | X | X |
| cysteine protease staphopain A | X | X | X | X |
| alpha-hemolysin | X | X | X | X |
| bi-component gamma-hemolysin HlgAB subunit A |  |  | X | X |
| bi-component gamma-hemolysin HlgAB/HlgCB subunit B | X | X | X | X |
| bi-component gamma-hemolysin HlgCB subunit C | X | X | X |  |
| staphylococcal enterotoxin type A |  | X |  |  |
| staphylococcal enterotoxin type R |  | X |  |  |
| staphylococcal protein A |  |  |  | X |
| bi-component leukocidin LukGH subunit G | X | X | X | X |
| bi-component leukocidin LukGH subunit H | X | X | X | X |
| bifunctional autolysin | X | X | X | X |
| phenol-soluble modulins PSM-alpha-4 | X | X | X |  |
| phenol-soluble modulins PSM-alpha-x |  |  | X |  |
| beta-class phenol-soluble modulins | X | X | X |  |
| delta-lysin family phenol-soluble modulins | X |  | X |  |
| serine protease SplB | X | X |  |  |
| serine protease SplD |  | X |  |  |
| serine protease SplE |  | X |  |  |
| serine protease SplF |  | X |  |  |
| Fur family transcriptional regulator |  | X |  |  |
| staphylobilin-forming heme oxygenase IsdI | X | X | X | X |
| LPXTG-anchored heme-scavenging protein IsdA | X | X | X |  |
| von Willebrand factor binding protein Vwb |  | X | X | X |
| two-component system response regulator SrrA | X | X | X | X |
| two-component system sensor histidine kinase SaeS | X | X | X | X |
| HTH-type transcriptional regulator SarS |  |  | X | X |

Bacterial uptake by U937 cells was verified using transmission electron microscopy (TEM; **Fig. 1 Ai-Aiv**) and imaging flow cytometry (**Fig. 1 Av-Aviii**), and by SaOS-2 and A549 cells (**Fig. 1Bi-Cii**). At 2 hpi, *S. aureus* was internalized and primarily resided within membrane-enclosed compartments of varying morphology (size, wide or tightly packed, reflecting different maturation states^39^), and varying numbers of engulfed bacteria. Most vacuoles contained replicating bacteria, whereas degrading *S. aureus* was found particularly in U937 cells (e.g. **Fig. 1 Ai**, blue arrow). A notable fraction of bacteria localized perinuclearly or intranuclearly in SaOS⍰2 (EDCC 5055 **Fig. 1 Bi, Biii**, EDCC 5464 **Fig. 1 Bii, Biv**), A549 cells, and less frequent in U937 cells (**Fig. S3**). SBF-SEM confirmed intranuclear localization in SaOS-2 (**Fig. 1 D-F**). SIM imaging further demonstrated colocalization with the nuclear surface for EDCC 5055 (**Fig. 1G**) and EDCC 5464 (**Fig. 1H**) (SH1000, EDCC 5458, and imaging of A549 cells, see **Fig. S3**) at distinct relative frequencies (**Fig. 1I**). LAMP-1 proximity (**Fig. 1J**) suggested partial bacterial residence within lysosomal compartments at 2 hpi, dependent on each isolate.

**Fig. 1.**
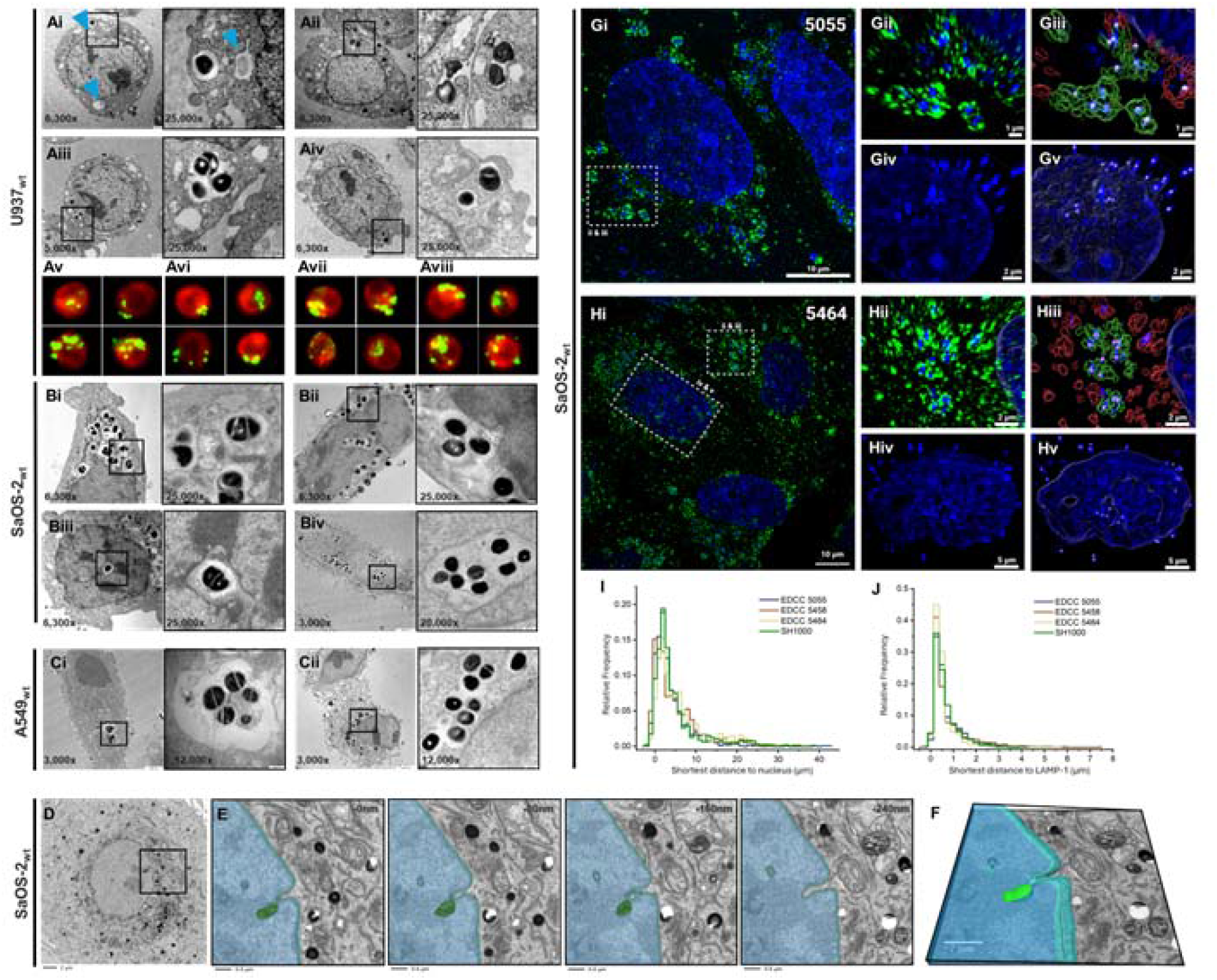
*S. aureus* isolates reside intracellularly in LAMP-1 associated compartments, in the cytosol as and in the host nuclei. Each cell line was infected with *S. aureus* at MOI 100 using a gentamicin protection assay and cells were fixed at 2 hpi. Transmission electron microscopy of (**A**) U937, (**B**) SaOS-2, and (**C**) A549 cells infected with two or four of the *S. aureus* isolates (**Ai**) SH1000, (**Aii, Bi, Biii, Ci**) EDCC 5055, (**Aiii**) EDCC 5458, and (**Aiv, Bii, Biv, Cii**) EDCC 5464. Representative images at two different magnifications (black box, 3,000x-85,000x as indicated) for n=1 (U937) or n=2 (A549, SaOS-2) independent experiments, each with n≥10 analyzed cells. Blue arrows indicate possible degrading bacteria. (**Av-viii**) imaging flow cytometry analysis shows CFSE-labelled bacteria in green inside U937 host cells (red; CellMask deep red). (**Av**) SH1000, (**Avi**) EDCC 5055, (**Avii**) EDCC 5458, (**Aviii**) EDCC 5464. (**D**) SBF-SEM analysis of EDCC 5055-infected SaOS-2 cells shows intranuclear localization. (**E**) Zoom-in of boxed region in D as pseudo-coloured sections of a total of 240nm and (**F**) their 3D-reconstruction. (**G, H**) Maximum intensity projection of SaOS-2 cells infected with (**Gi**) EDCC 5055 or (**Hi**) EDCC 5464 imaged by SIM. DNA is shown in blue, Lamp-1 positive organelles in green. (**Gii, Hii**) Zoom-in of boxed region in Gi/Hi. (**Giii, Hiii**) Surfaces and spots generated after image analysis in Imaris (11.0.0). Surfaces defining the nucleus are visualized by a blue outline, spots corresponding to bacterial DNA are highlighted by white spheres. Surfaces generated around LAMP1 signal are classified based on distance to the closest bacterial spot. Surfaces closer than 0.25 µm coloured in green, while surfaces at a greater distance are coloured in red. (**Giv, Hiv**) DAPI stained nucleus of a cell infected with EDCC 5055/5464 (not depicted in Gi). Intensive DAPI staining consistent with bacterial DNA stain can be observed inside the nucleus or located within an invagination of the nuclear lamina. (**Gv, Hv**). Spots are identified inside the generated nucleus surface. (**I**) Histogram of shortest distances between identified bacterial spots and the surface generated around the nucleus (bin size 1 µm; 8 images per strain; 1515 ± 160 (mean ± SD) detected spots per strain). (**J**) Histogram of shortest distances between identified bacteria spots and the surface generated around LAMP1 signals (bin size 0.25 µm; 8 images per strain; 1515 ± 160 (mean ± SD) detected spots per strain).

### *S. aureus* infection induces a conserved transcriptional stress signature across isolates alongside a distinct, isolate-specific response in U937 cells

Circulating and tissue-resident monocytes and macrophages are among the first line of defence countering invasive *S. aureus*. Hence, we performed RNA sequencing of U937 cells at 4 hpi (MOI 30) to determine their transcriptional response. Principal component analysis (PCA) (**Fig. 2A**) revealed pathogen-specific host responses: *S. aureus* infection resulted in variable clustering depending on the isolate, whereas *E. faecium* infection, included as gram-positive control bacterium, was less variable. Differentially expressed gene (DEG) analysis comparing *S. aureus* infection with the uninfected control (**Fig. 2B-E**) underlined a divergent but strong transcriptional response, while several of the top up- and downregulated genes were conserved between isolates (for interactive volcano plots, see **Supplementary Files S4-S9**). To identify functionally relevant expression patterns, we selected only protein-coding significant DEGs (padj < 0.05, |log2FC| ≥ 1). Further analysis is shown here for *S. aureus*-infected cells only.

**Fig. 2.**
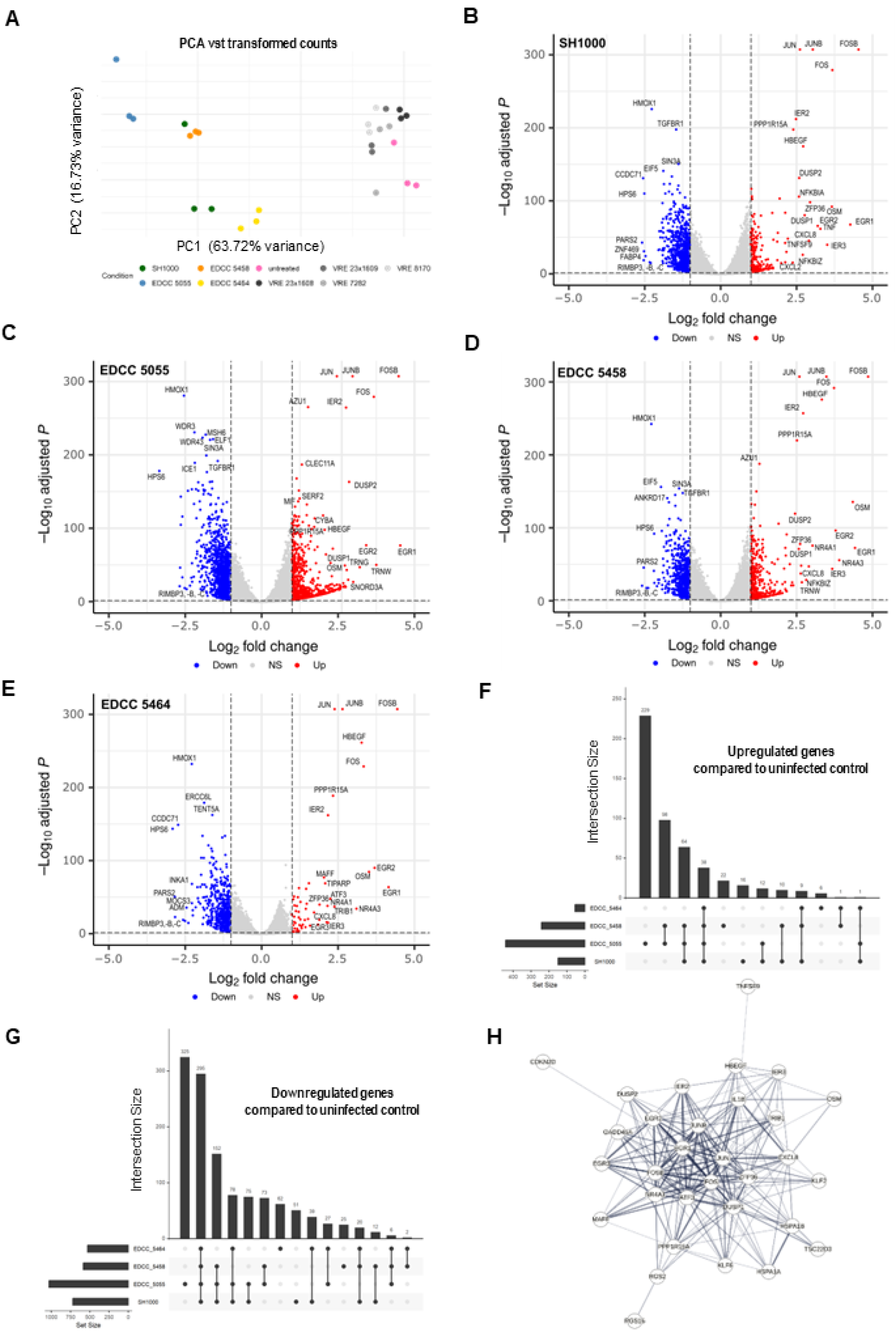
U937 cells infected with *S. aureus* reveal both shared and isolate-specific differentially regulated gene signatures. U937 cells were infected in triplicates with different *S. aureus* (4 hpi) using a gentamicin protection assay at MOI 30 or, for comparison, with *E. faecium* (6 hpi) isolates. The host’s transcriptional response was analyzed using Illumina mRNA sequencing. **(A)** Principal component analysis (PCA) of variance-stabilizing transformation (vst)-transformed data using the top 1000 most variable genes shows close clustering of VREfm-infected, but scattered clustering for *S. aureus*-infected cells compared to untreated controls. Further analyses are shown for *S. aureus* only. Volcano plots of cells infected with *S. aureus* isolates (**B**) SH1000, (**C**) EDCC 5055, (**D**) EDCC 5458, and (**E**) EDCC 5464 compared to uninfected control cells delineate overlapping and unique transcriptional regulation (padj < 0.05, |Log2FC| >= 1), supported by UpSet plots depicting overlap of significantly, at least two-fold (**F**) upregulated and (**G**) downregulated genes across conditions. **(H)** Full STRING network of shared upregulated, protein-coding genes (padj < 0.05, |Log2FC| ≥1) across all *S. aureus* isolates (edges indicate confidence, medium confidence 0.400), disconnected nodes not shown.

The overall number of DEGs as well as the number of shared and unique upregulated (**Fig. 2F**) and downregulated (**Fig. 2G**) genes varied strongly between infection with the different isolates, peaking for EDCC 5055, and was lowest for EDCC 5464 exposed cells. Overall, 295 genes were downregulated by all four isolates, while those upregulated covered 38 genes (**Fig. 2H**). These were among those of the highest relative expression and are in line with an isolate-independent stress response, transcriptionally regulating signal transduction, cell survival, and proliferation (e.g. *FOS, FOSB, JUN, JUNB, EGR1/2, IER2/3, HSPA1A/B, ZFP36*), as well as inflammation (*CXCL8, IL1B*).

To functionally analyze shared (**Fig. 3A**) and unique (**Fig. 3B**) modulated pathways in infected versus uninfected control cells, we performed GO:BP pathway enrichment analysis and summarized the results via Jaccard hierarchical clustering (*h=0*.*5*) of significant gene sets. Directly comparing expression between cells infected with different isolates underlined distinct host response patterns (**Fig. 3C**, representatively for SH1000 vs. EDCC 5055 and EDCC 5055 vs. 5464). Shared clusters still markedly varied in their respective enrichment score and the number of involved genes across isolates.

**Fig. 3.**
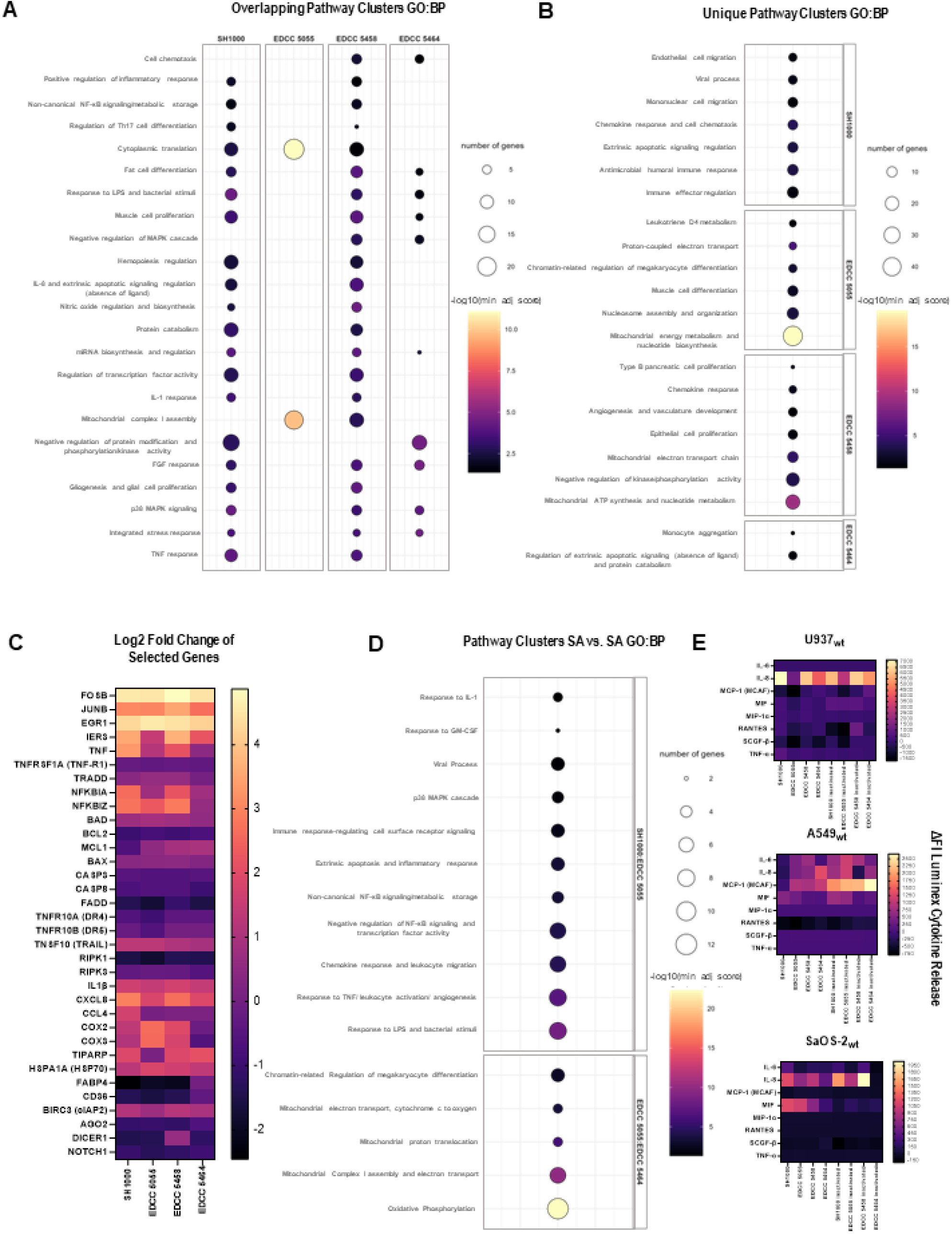
Intracellular *S. aureus* infection of U937 cells induces distinct upregulated pathway clusters dependent on each isolate as well as cell line- and isolate-specific cytokine release patterns. Significantly (padj≤0.05), at least two-fold upregulated protein-coding genes were analyzed for enriched pathways (Gene Ontology Biological Processes) using g:Profiler ordered query, and clustered based on Jaccard similarity of gene sets, using hierarchical clustering (average linkage) and a fixed cut height of 0.5. The adjusted enrichment score of a driver pathway (minimal padj) is shown representatively for each cluster (-log10(min_adj_score)) and bubble size indicates gene set size, for (**A**) overlapping and (**B**) unique pathway clusters of *S. aureus* isolates compared to untreated controls. (**C**) Log2 fold changes of 20 selected genes of interest against untreated control. (**D**) overlapping and unique clusters directly comparing two isolates (SH1000 vs. EDCC5055 and EDCC 5055 vs. EDCC 5464). (**E**) Median fluorescence intensities normalized to uninfected control cells and background (ΔFI) of eight selected cytokines using Luminex for U937, A549 (n=3 for viable, n=2 for inactivated *S. aureus*), and SaOS-2 (n=1) cells.

The isolates SH1000, EDCC 5458, and EDCC 5464, but not EDCC 5055, strongly induced negative regulation of kinase and phosphorylation activity, involving e.g. *CDKN1A/2D, DUSP1 or GADD45*. While EDCC 5055 induced the highest number of individual DEGs, these resulted in more distinct clusters compared to the other isolates. Upregulated pathways were primarily linked to mitochondrial respiration and energy metabolism (e.g. *COX1, -2, -3, CYTB, MT-ND1, -2 or ATP6, -8*), some of which were uniquely induced by this isolate (*MT-ND1, COX1*). Mitochondrial pathways were also upregulated for EDCC 5458 to a lesser extent in comparison to EDCC 5055, but not upon SH1000 or EDCC 5464 infection (e.g. *COX1* is downregulated by the latter). EDCC 5055 further upregulated cytoplasmic translation (ribosomal proteins of the 40S and 60S subunits, e.g. *RPL24, -26, -31, RPLP1*, OR *RPS12, -14, -15*), chromatin-organization associated pathways (e.g. *H4C1, H4C5, H4C11, H4C12, H4C14, H4C15 or H1-2, H1-4*). Interestingly, leukotriene D4 metabolism (unique upregulation of e.g. DPEP2 or GGT5) represented the only inflammation-associated (single) pathway cluster for this isolate.

In contrast, the isolates EDCC 5458 and, particularly, SH1000 induced broader immune response- and inflammation-related (e.g. monocyte activation marker CD69 (SH1000), chemokines and cytokines *CCL20, CCL4* (unique for SH1000), *CXCL2, CXCL8, IL1B, TNF, NFKBIA, NFKBIZ, OSM*), and apoptosis-regulating pathway activation (e.g. NF-κB-induced *BIRC3, CIAP2, TNF, NFKBIA, NFKBIZ*). The latter are critically related to TNF signaling (log2FC of 3.3 and 2.1 for SH1000 and EDCC 5458, respectively, of TNF). Notably, marked differential TNF expression and TNF-related pathway responses comparatively lacked upon infection with EDCC 5055 or 5464. TNF-induced protein 3 (*TNFAIP3*) was uniquely upregulated upon SH1000 infection, whereas its expression was uniquely downregulated for EDCC 5055. RIPK1, but not RIPK3, was significantly downregulated by all isolates. For SH1000, EDCC5458, and EDCC5464, immune and apoptosis-related pathways clusters were enriched, driven by genes involving e.g. *CXCL8, IL1B, HSPA1A/B* (encoding HSP70), TIPARP (primarily involved in antiviral signaling and not induced by EDCC 5055), or *PP1R15A* (related to growth arrest and autophagy, lysosomal biogenesis), as well as MAPK-related genes (e.g. *DUSP1, RGS2, GADD45A/B*). Positive and negative regulation of MAPK-related cascades was associated with SH1000 and EDCC 5464, respectively, while both were upregulated by EDCC 5458. On the other hand, EDCC 5055 infection did not result in enrichment of these pathway clusters, despite differential regulation of single related genes.

Expression of several key regulators of apoptosis was significantly modulated by some isolates compared to uninfected control cells, but they were below or near the cut off. These include anti-apoptotic signaling genes, such as *BAD, BCL2* (for SH1000 over cut off), or *MCL1*. Except e.g. *BAX*, many pro-apoptotic genes were downregulated, including caspase-3 and -8, associated with moderate (over-cut-off) downregulation of *FADD* across all isolates and attenuated death receptor expression, e.g. *TNFRSF1A* (TNF-R1), *TNFR10A* (DR4) or *TNFR10B* (DR5; only for SH1000 and EDCC 5055). TRADD (not for EDCC 5464) and *TNSF10* (TRAIL) were moderately upregulated for all isolates below or near the cut-off. Neither CD95/Fas, nor FasL were differentially regulated by any of the isolates. Log2FC values of selected genes of interest compared to the uninfected control that were significantly regulated for at least one isolate are shown in **Fig. 3C**.

These isolate-specific functional signatures became particularly apparent upon directly comparing infected cells with different isolates, as shown for SH1000 vs. EDCC 5055 and EDCC 5055 vs. EDCC 5464 (**Fig. 3D**). All pathway clusters upregulated by SH1000 compared to EDCC 5055 were related to immune responses, inflammation, and programmed cell death regulation. Upregulated pathway clusters by the high-response isolate EDCC 5055 compared to low-response isolate EDCC 5464 were, except for chromatin-related regulation, entirely related to mitochondrial respiration.

Strikingly, EDCC 5464 failed to induce substantial specific transcriptional upregulation compared to the other isolates. Comparing DEGs of the other isolates to EDCC 5464 revealed that particularly genes related to fatty acid metabolism and transport (*FABP4, CD36*) as well as RNA stability and catabolic processes involved genes (e.g., *AGO2, DICER1, DHX36*) were significantly downregulated in all isolates but the latter (**Supplementary Fig. S10**). Other downregulated pathway clusters covered Notch activation (e.g., *NOTCH1, MAML1-3*) for SH1000 and EDCC 5458, while EDCC 5055 downregulated pathways related to p53-mediated (e.g., *MYC*) or TGF-β (e.g., *TGFBR1*) signaling, epigenetic regulation of gene expression (e.g. *ALKBH1/-4*), small GTPase signaling (e.g. *ABL2*), or ribosome biogenesis (e.g. *DDX28*), amongst others. The full lists of DEseq2 data, overall significant, unique and shared DEGs as well as enriched pathways, clusters, and included genes are available in **Supplementary File S11**.

Together, our findings reveal both a strong transcriptional signature of U937 cells in response to intracellular infection that is conserved across all tested *S. aureus* isolates as well as unique transcriptional regulation specific to each isolate. Differentially affected pathways are primarily related to mitochondrial respiration (EDCC 5055), (TNF-associated) inflammation (SH1000, EDCC 5458), and regulation of apoptosis (SH1000, EDCC 5458, EDCC 5464). Notably, the isolate of a chronic infection background, EDCC 5464, barely induces specific transcriptional upregulation in U937 cells.

### Isolate- and cell line-dependent cytokine release that is not restricted to viable *S. aureus*

We next screened for cytokine release, initially using Luminex 48-plex ELISA in infected U937 cells and A549wt cells (**Supplementary Fig. S12**), then stratifying to eight cytokines for further testing additionally in SaOS-2 cells at 4 hpi, employing both viable and heat-inactivated *S. aureus* (**Fig. 3E**). Analysis revealed distinct cytokine release patterns, underscoring that both host cell type, bacterial isolate, and bacterial viability shape host cytokine induction.

Compared to their uninfected controls, all cell lines showed IL-8 release at varying levels and to an isolate-dependent extent, and was most pronounced in U937 cells. A549 preferentially released MCP-1 (MCAF) and dampened RANTES secretion. IL-6 was released from A549 and to a lesser extent from SaOS-2 cells, while completely lacking in U937 cells, mirroring absent transcriptomic regulation of this cytokine. Heat-inactivated isolates still stimulated cytokine release to a comparable level or even exceeding that of viable bacteria, supporting that viable *S. aureus* actively modulates host immune signaling, which varies between cell lines. Whereas inactivated EDCC 5458 markedly increased IL-8 in U937 and SaOS-2 cells, release was reduced below control levels in A549 cells. SaOS-2 cells selectively released MIF in response to viable isolates. Conversely, U937 cells released MIF only upon treatment with heat-inactivated, but not with viable *S. aureus*. TNF-α secretion showed marked isolate- and host cell dependency and was predominantly induced by U937 cells infected with SH1000 and, to a lesser extent, EDCC 5458 or 5464, but was mainly absent with EDCC 5055 or in A549 and SaOS-2 cells across isolates. Indeed, overall cytokine release from U937 cells was attenuated upon infection with viable as well as heat-inactivated EDCC 5055 compared to the other isolates, while overall fluorescence intensities peaked in U937 cells versus non-professional phagocytic cell lines.

### Intracellular *S. aureus* triggers U937 host cell death to an isolate-dependent extent, involving several cell death pathways

We next aimed to elucidate the host cell fate by analyzing infection-induced cell death in U937, compared to A549 and SaOS-2 cells.

In U937 cells, Annexin V/ 7-AAD staining revealed host cell death signaling by all *S. aureus* isolates as early as 6 hpi, peaking upon challenge with EDCC 5055 (**Fig. 4A, left**). Notably, preventing direct bacteria-host cell contact (bc: boyden chamber), cell death is abrogated entirely at 6 hpi, emphasizing the requirement for contact or uptake (**Fig. 4A, right**). Without bc, isolate-specific host cell death levels did not substantially increase from 6 hpi to 24 hpi (**Fig. 4B, left**). However, upon bc application, cell death induction is restored at 24 hpi, significantly for EDCC 5055 and by trend for EDCC 5458 and 5464 (**Fig. 4B, right**). Significant activity of the pro-apoptotic caspases-3 or -7 was observed only for EDCC 5055 at 6 hpi (**Fig. 4C, Fig. 4E**) and then barely further increased until 24 hpi for all isolates (**Fig. 4D, Fig 4F)**, suggesting isolate-dependent apoptosis activation during earlier stages of infection.

**Fig. 4.**
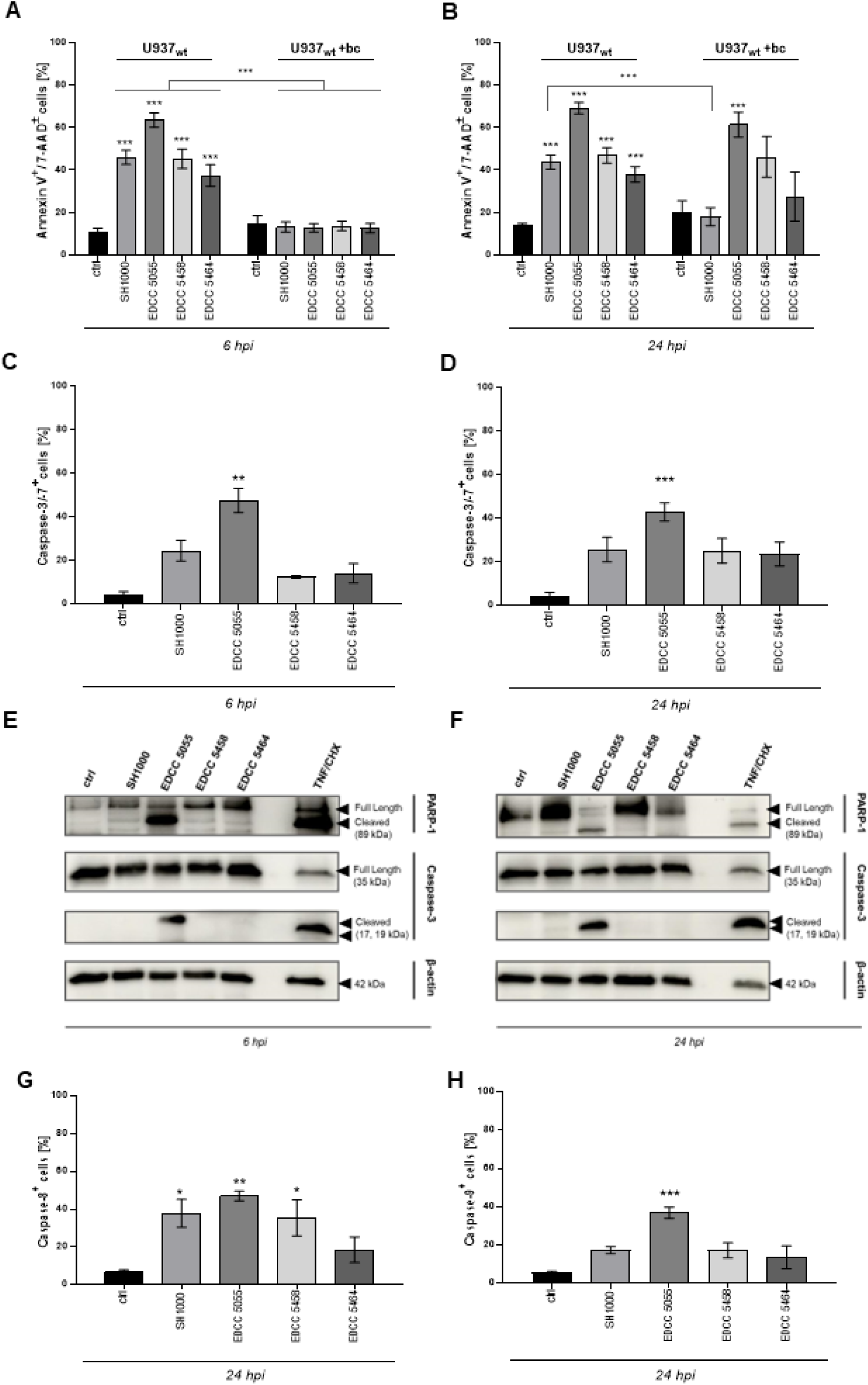
Intracellular *S. aureus* induces cell death of U937 cells to an isolate-dependent extent with apoptotic effector activity at earlier stages of infection. Cells were infected with *S. aureus* isolates at MOI 30 using a gentamicin protection assay or direct contact being prevented using cell culture inserts (Boyden Chamber, +bc). (**A, B**) At 6 hpi and 24 hpi AnnexinV^+^/ 7-AAD^±^ cells, (**C, D**) Caspase-3/-7^+^ at 6 hpi and 24 hpi. Protein expression of the full-length and cleaved form of PARP-1 and Caspase-3 was confirmed via Western Blot at (**E**) 6 hpi and (**F**) 24 hpi. (**G**) Caspase-8^+^ and (H) Caspase-9^+^ cells at 24 hpi were quantified using flow cytometry. Data show mean ± SEM of n≥3 independent experiments. Cells were sensitized with cycloheximide (CHX) (0.5 µM) 30 min prior to treatment with TNF (100 ng/mL) as a positive control. Statistical analysis was performed using Two-way ANOVA with Tukey’s Multiple Comparisons between all conditions. Significance is shown for each *S. aureus* isolate versus uninfected control as well as for the same isolate between groups (with and without bc; indicated by lines); not shown for isolate versus isolate within the same group. *p<0.05; **p<0.01; ***p<0.001.

Despite only marginal caspase-3/-7 effector activity induced by the other isolates, all isolates except EDCC 5464 elicited significant activation of the initiator caspase-8 at 24 hpi (**Fig. 4G**), indicating upstream activation of the extrinsic apoptotic pathway. Significant caspase-9 activity was observed exclusively in cells exposed to EDCC 5055 (**Fig. 4H**), suggesting additional intrinsic apoptosis. However, a proportion of cells lacked activity of these pro-apoptotic caspases, but were Annexin V and/or 7-AAD positive, and pan-caspase inhibition using zVAD.fmk did not prevent cell death (not shown). Because both necroptosis and pyroptosis permit 7-AAD uptake and Annexin V also marks pyroptotic cells ^40^, we examined further effectors not specific to apoptosis.

### TNF-R1 dictates the timing and extent of isolate-specific host cell death in response to intracellular *S. aureus* in U937 monocytes, but not A549 or SaOS-2 cells

We previously observed isolate-specific TNF transcriptional regulation, cytokine release, and activation of TNF-related pro-inflammatory and survival pathways in U937 cells, suggesting TNF-R1 to modulate cell fates upon infection. Strikingly, *S. aureus*-induced host cell death (Annexin V^+^/7-AAD^±^) was completely abrogated in U937 cells lacking TNF-R1 (U937_ΔTNF-R1_) at 6 hpi (**Fig. 5A**), but was restored at 24 hpi, however at significantly different levels compared to U937_wt_ cells for each isolate (**Fig. 5B**). Given the potential involvement of apoptotic and non-apoptotic signaling, simultaneous multi-caspase profiling revealed isolate-specific activity that exceeded caspase-3/-7, -8, or -9 alone at 2411hpi (**Fig.⍰5C**), while multi-caspase and caspase-3/-7 activity were comparable at 6⍰hpi (**Supplementary Fig.⍰S13**).

**Fig. 5.**
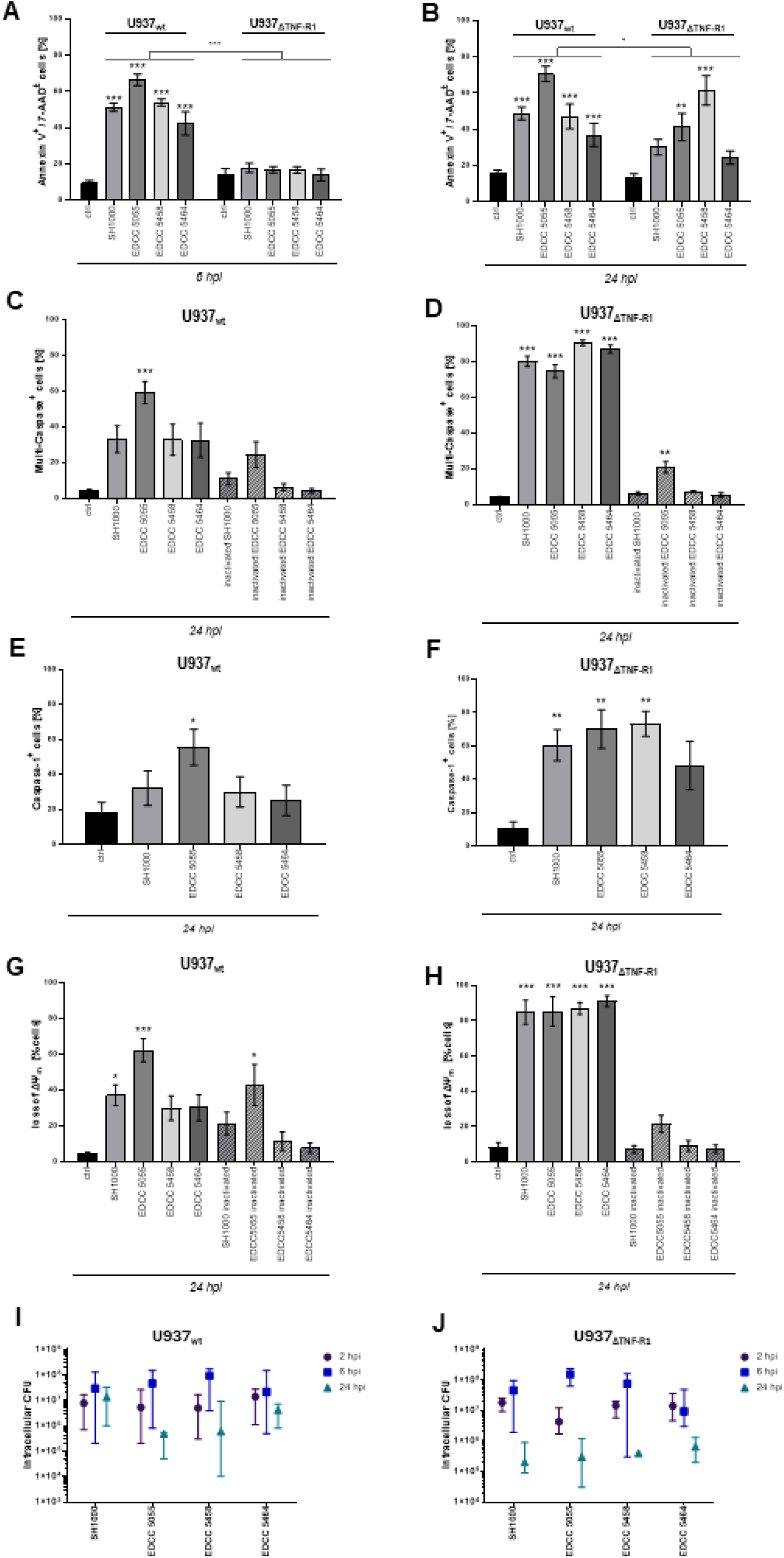
TNF-R1 signaling temporally controls isolate-specific U937 cell death during intracellular *S. aureus* infection. U937 wildtype and U937_ΔTNF-R1_ cells were infected with viable or heat-inactivated *S. aureus* isolates at MOI 30 using a gentamicin protection assay. (**A**) At 6 hpi and (**B**) 24 hpi, Annexin V^+^ / 7-AAD^±^ cells (Dead Cells) were analyzed using flow cytometry. At 24 hpi, Multi-Caspase ^+^, Caspase-1^+^ (FAM-YVAD-FMK), and depolarized (loss of mitochondrial transmembrane potential, ΔΨm) cells were quantified by flow cytometry in (**C, E, G**) U937_wt_ and (**D, F H**) U937_ΔTNF-R1_ cells, respectively. Data show (**A-H**) mean ± SEM of n≥3 independent experiments. (**I**) U937_wt_ and (**J**) U937_ΔTNF-R1_ cells were lysed at the indicated time-points, and intracellular colony-forming units (CFU) were quantified per lysate (initial concentration: 1×10 cells). Data show median ± 95% CI on log-scale, zero-values were excluded. Statistical analysis was performed using One- or Two-way ANOVA with Tukey’s or Kruskal-Wallis Multiple Comparisons comparing means between all conditions. Significance is shown for each *S. aureus* isolate versus uninfected control as well as for the same isolate between groups (**wt** versus **ΔTNF-R1**; indicated by lines); not shown for isolate versus isolate within the same group. *p<0.05; **p<0.01;***p<0.001.

Notably, excessive multiple-caspase activation was induced by all isolates, even EDCC 5464, in U937_ΔTNF-R1_ cells at 24 hpi and isolate-specific differences diminished (**Fig. 5D**). The involvement of non-apoptotic pathways was further supported by caspase-1 activity at levels comparable to multiple-caspases at 24 hpi in U937_wt_ (**Fig. 5E**), suggesting engagement of pyroptotic cell death at later stages of infection. Caspase-1 activation levels were again markedly amplified in U937_ΔTNF-R1_ cells.

U937 cells underwent significant changes in the mitochondrial transmembrane potential (ΔΨm), which is linked to several cell death modalities, in response to SH1000 and EDCC 5055 (**Fig. 5G**). This was equally potentiated in TNF-R1-deficient cells for all isolates. In both U937_wt_ and U937_ΔTNF-R1_ cells, several of these effects were restricted to viable *S. aureus* for all isolates but EDCC 5055.

The observed discrepancy between U937 phenotypes might be attributed to differential uptake efficacy. Quantification of intracellular CFU per lysate revealed comparable uptake rates in both wild-type (**Fig. 5I**) and TNF-R1-deficient (**Fig. 5J**) cells. CFU slightly increased from 2 to 6 hpi, hinting towards bacterial replication, followed by a sharp decline at 24 hpi for most isolates. While the latter might indicate bacterial degradation, it aligned with the observed cell death induction and thus bacterial escape, particularly for the isolates SH1000 and EDCC5464, which provoked the disintegration of only U937_ΔTNF-R1_ cells. Supporting this, we observed no quantifiable differences between isolates e.g. in bacterial fluorescence intensity or spot counts using imaging flow cytometry analyses (**Fig. 1 Av-viii**).

To assess *S. aureus*-induced cell death in the non-phagocytic cell lines A549 and SaOS-2, RTCA revealed cell detachment as a surrogate for cell death of A549 (**Fig. 6A**) and SaOS-2 (**Fig. 6B**) at isolate-dependent kinetics and magnitudes. EDCC 5055 and EDCC 5458 induced rapid detachment in both A549 and SaOS-2 starting at approximately 5 hpi. SH1000 concurrently initiated cell detachment, which then proceeded slower. In contrast, EDCC 5464-induced detachment was delayed by approximately 24 h and proceeded slowly within 48 hpi in A549 cells, while lacking entirely in SaOS-2 cells up to 72 hpi. In the latter, cell proliferation was however stagnant compared to uninfected controls. These observations are supported by intracellular bacterial loads of all isolates except EDCC 5464 declining in both A549 (**Fig. 6C**) and SaOS-2 (**Fig. 6D**), as shown by CFU per host cell lysate over 24 and 48 hpi, respectively.

**Fig. 6.**
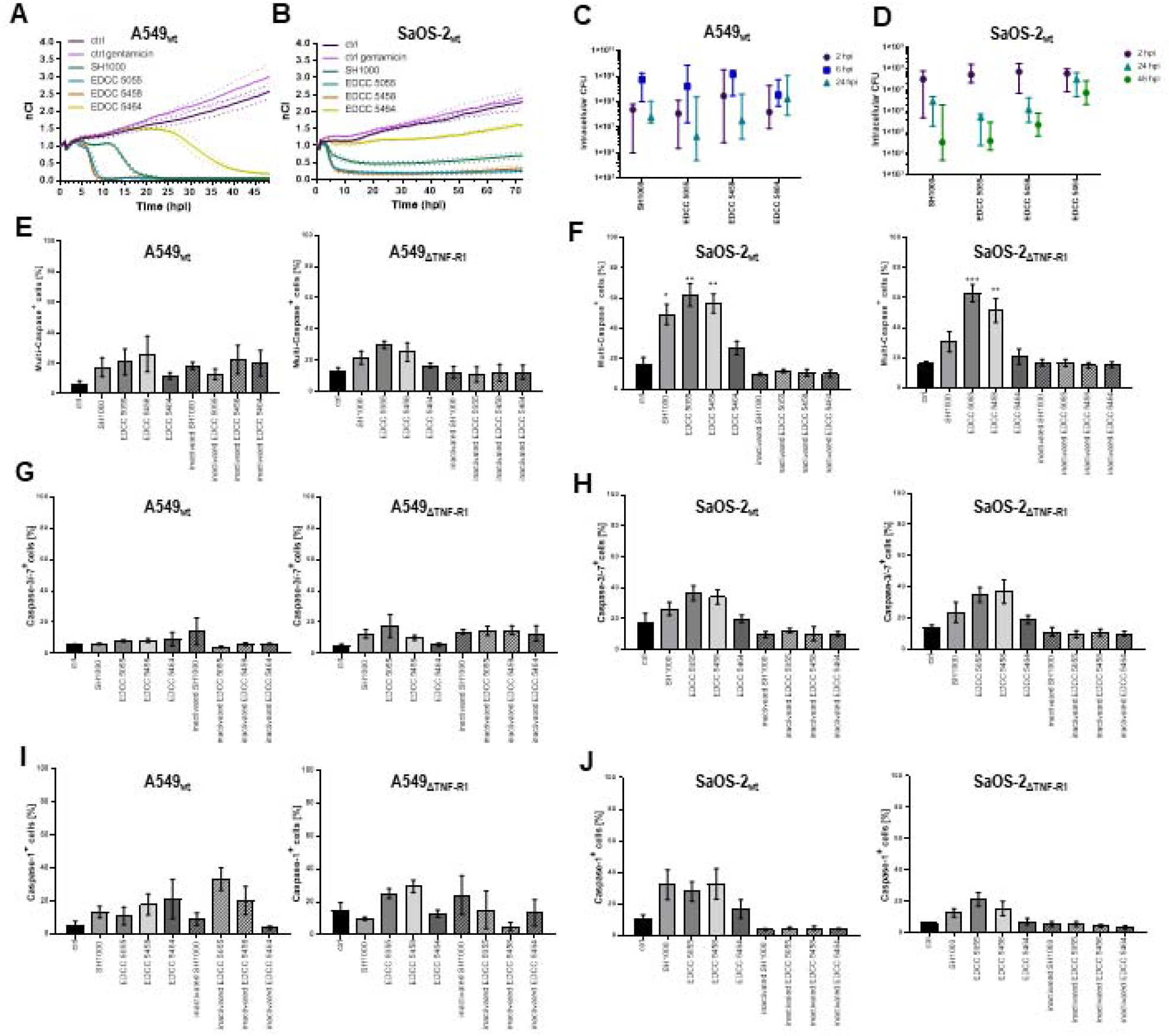
A549 and SaOS-2 cell death during intracellular *S. aureus* infection involves activation of various pathways to an isolate-specific extent, which is only moderately modulated by TNF-R1 signaling. A549_wt_ A549_ΔTNF-R1_, SaOS-2_wt_, and SaOS-2_ΔTNF-R1_ cells were infected with viable or heat-inactivated *S. aureus* isolates at MOI 30 using a gentamicin protection assay. Cell death kinetics of infected, untreated, and gentamicin-treated control **(A)** A549_wt_ and **(B)** SaOS-2_wt_ cells were normalized to the time of infection using xCELLigence RTCA. Detachment of cells indicating cell death is depicted as decreasing normalized Cell Index (nCI); one representative experiment is shown as mean ± SD (dotted line). At the indicated time-points, **(C)** A549_wt_ and (**D**) SaOS-2_wt_ cells were lysed, and intracellular colony-forming units (CFU) were quantified per lysate (initial concentration: 5×10^5^ cells). Data show Median ± 95% CI on log-scale, zero-values were excluded. At 24 hpi, Multi-Caspase ^+^, Caspase-3/-7 ^+^, and Caspase-1^+^ cells were analyzed by flow cytometry in (**E, G, I**) A549_wt_ (**left**) and A549_ΔTNF-R1_ (**right**) and **(F, H, J)** SaOS-2_wt_ and (**left**) SaOS-2_ΔTNF-R1_ (**right**) cells, respectively. Data show mean ± SEM of n≥3 independent experiments. Statistical analysis was performed using One-way ANOVA with Tukey’s or Kruskal-Wallis Multiple Comparisons. *p<0.05; **p<0.01; ***p<0.001 vs. uninfected controls.

Next, we analyzed caspase activation and impact of TNF-R1 in these cell lines. In A549_wt_ cells, activity of multiple caspases (**Fig. 6E, left**) as well as caspase-1 (**Fig, 6I, left**) was attenuated, while *S. aureus* failed to trigger caspase-3/-7 activation (**Fig. 6G, left**) at 24 hpi. Knock-down of TNF-R1 only slightly reinforced multi-caspase (**Fig. 6E, right**), caspase-3/-7 (**Fig. 6G, right**), and caspase-1 activities (**Fig. 6I, right**). In both A549_wt_ and, for caspase-1, in A549_ΔTNF-R1_, also heat-inactivated *S. aureus* showed effector-dependent stimulation, partially exceeding that of their viable counterparts.

In SaOS-2 cells, significant activation of multiple caspases (**Fig. 6F**) was induced by viable SH1000, EDCC 5055, and EDCC 5458 in wild-type (**Fig. 6F, left**) and only EDCC 5055 and EDCC 5458 in TNF-R1 deficient (**Fig. 6F, right**) cells at 24 hpi. In contrast to U937 cells, caspase-3/-7 and -1 were activated with only minor isolate-specific differences in either phenotype, whereas the latter was slightly attenuated in SaOS-2_ΔTNF-R1_ compared to SaOS-2_wt_. TNF-R1 deficiency retained isolate-specific effects in both A549 and SaOS-2 cells, albeit being less pronounced in the first place.

Altogether, these findings show cell line- and isolate-dependent host cell fates during intracellular *S. aureus* infection, with heterogenous caspase-dependent death particularly in U937 and SaOS-2 cells. TNF-R1 knock-down uncovers its biphasic role in U937 monocytes, delaying initial regulated cell death to late, excessive cell destruction driven by broad effector activation. As these effects require viable bacteria (except EDCC 5055), the isolates likely manipulate TNF-R1 signaling in distinct, cell type-specific ways to modulate host cell death versus survival.

## Discussion

Intracellular infection of various professional and non-professional phagocytic host cell types is now a well-studied feature of *S. aureus*. However, many studies focus on host cell modulation in single cell lines induced by an individual isolate, neglecting the interplay of specific isolate- and tissue type-dependent properties to shape cellular outcome ^6,9^. In this study, we demonstrate that intracellular *S. aureus* elicits a conserved early stress response across four isolates yet triggers highly isolate-specific transcriptional as well as both isolate- and cell type-specific cytokine and death programs. In U937 monocytes, TNF-R1 emerges as a central gatekeeper balancing regulated versus uncontrolled cell death in an isolate-dependent manner, whereas its contribution is far less pronounced in A549 and SaOS-2 cells. Finally, our findings suggest peri- and intranuclear localization across cell types as a previously unidentified intracellular niche.

### Intracellular residence with peri- and intranuclear localization across isolates

In line with previous studies ^6,9,41^, we show that all analyzed isolates are rapidly internalized by all cell lines, primarily residing in membrane-enclosed compartments in mostly replicating but also degrading states at 2 hpi. These compartments show LAMP-1 decoration and commonly cluster perinuclearly, consistent with the known spatial organization of endo-lysosomes promoting membrane fusion and autophagic flux ^42^. However, many pathogens, including *S. aureus*, are known to interfere with lysosomal trafficking and fusion ^10^. Perinuclear lysosome accumulation in *S. aureus*-infected macrophages has been linked to the small GTPase Arl8 ^43^ and becomes apparent in other TEM studies of *S. aureus*-infected host cells, demonstrating bacteria-containing invaginations of the nuclear membrane or even empty vacuoles within the nucleus ^44^. Similar localization has been reported for a few bacterial species, including the human pathogens *O. tsutsugamushi* or *R. rickettsii* ^45^. Notably, our findings reveal also *intranuclear* bacterial localization either within membrane-bound vacuoles or directly in the nucleoplasm at 2 hpi, raising the possibility that *S. aureus* accesses and exploits nuclear compartments through manipulation of vesicle trafficking or nuclear envelope integrity. Further spatio-temporal analyses will be required to determine whether this non-canonical localization represents a transient trafficking stage or a distinct niche supporting persistence or immune evasion.

### Isolate-specific transcriptional programs of U937 monocytes in response to S. aureus

Transcriptomic profiling of U937 monocytes at 4 hpi reveals a robust stress response (e.g. FOS, JUN, OSM, CXCL8, IL1B) ^46^ conserved across isolates, and a highly divergent, isolate-specific transcriptional signature. The strains SH1000, EDCC 5055, and EDCC 5458 elicit hundreds of DEGs, as opposed to the muted response to the chronic infection isolate EDCC 5464. Each strain triggers unique pathway activation. For instance, EDCC 5055 most prominently upregulates mitochondrial energy metabolism and translational pathway cluster, while particularly SH1000 and also EDCC 5458 robustly provoke e.g. TNF, NF-κB, and MAPK-related inflammatory and apoptosis-related programs. In addition, both isolates moderately trigger TNF release. At least for SH1000, this observation is potentially linked to the unique upregulation of TNFAIP3, thus negatively regulating TNF expression in infected cells and possibly autocrine apoptosis ^47,48^, whereas it is uniquely downregulated by EDCC 5055. Interestingly, all isolates but EDCC 5464 downregulate FABP4 and CD36 transcription, linked to fatty acid metabolism, while the latter has also been implicated as a phagocytic receptor for *S. aureus*, with CD36-deficiency resulting in impaired TNF production and bacterial clearance ^49^.

Inflammatory signal transduction pathways are not transcriptionally enhanced upon infection with highly cytotoxic EDCC 5055, which might reflect anti-inflammatory and immune evasive strategies commonly employed by intracellular *S. aureus* to improve its own survival ^47,50-52^. Interestingly, genes related to leukotriene metabolism are upregulated (DPEP2, GGT5, GGTL1, GGTL2). In non-*S. aureus* inflammatory diseases, DPEP2 is reported to be gradually upregulated with macrophage differentiation, acting as a cell-type specific key regulator for macrophage-mediated inflammation by repressing NF-κB and MAPK activity as well as TNF, IL-6, or MCP-1 release ^53,54^. This is supported by transcriptional and cytokine-release responses observed in our study in the undifferentiated monocytes (see below). Pro-inflammatory cytokines are further associated with the expression of growth factors, which can provoke pathogenic fibrotic processes by affecting cellular differentiation. For instance, in bovine epithelial cells, *S. aureus* induces expression of the growth factors FGF and TGF-β via TLR-NF-κB and TLR-AP-1 (FOS/JUN) transcriptional activity ^55^, but also monocytes are reported to participate in the regulation of angiogenesis and endothelial function via growth factor signaling ^52^. In our model, all isolates but EDCC 5055 induce pathways implicated in response to FGF, whereas both EDCC 5055 and EDCC 5458 downregulate TGF-β receptor signaling.

All isolates suppress genes governing NLS-bearing nuclear import as well as RNA stability and catabolism regulation (e.g. AGO2, DICER1; except for EDCC 5464) compared to uninfected control cells. Dysregulated RNA stability may aid the pathogen, as Ago2-bound miRNAs modulate neutrophil bactericidal activity at *S. aureus*-infected wound sites ^56^. EDCC 5055 suppresses p53-linked signaling and epigenetic regulators while upregulating chromatin-organization genes, consistent with epigenetic dysregulation reported in intracellular *S. aureus* infection ^11,57^, which could hint to non-transcriptional regulation levels. Accordingly, whereas *S. aureus*-induced neutrophil lysis is shown to require de novo gene transcription and protein synthesis ^39^, key cellular pathways can also rely on transcription-independent regulation ^17^. Given the altered phosphorylation and protein-modification pathways in U937 cells, targeted organelle proteomic analyses, including PTMs, and profiling of non-coding RNAs such as miRNAs may help resolve key signaling mechanisms. *S. aureus*-induced transcriptional changes in non-professional phagocytes, such as osteoblasts ^11^ and endothelial cells ^7,47^, show both overlap and divergence from U937 responses observed here. Cell line-specific effects and comparison to primary cells will be essential to delineate isolate-specific host reactions.

### Isolate- and cell line-specific cytokine release upon S. aureus intracellular infection

Pro⍰inflammatory cytokine secretion is a key host defence against intracellular *S. aureus*. U937 and SaOS-2 cells release notable, and A549 moderate, levels of IL⍰8 with clear isolate- and cell-type specificity, highlighting tissue-dependent immune microenvironments. Notably, heat-inactivated isolates trigger cytokine levels comparable to or exceeding live bacteria, implying active suppression by viable *S. aureus* ^20,51^. *S. aureus* is frequently reported to induce IL-6 release with bacterial dose-dependent effects ^46,47,58^. Interestingly, an isolate-specific IL-6 response is observed mainly in A549, less pronounced in SaOS-2, and entirely absent in U937 cells in our models, with moderate TNF release only in the latter. Only osteoblast-like SaOS-2 selectively release MIF following infection with viable *S. aureus* (except EDCC 5464), consistent with studies reporting this cytokine to amplify pro-inflammatory and anti-proliferative signaling via NF-κB in *S. aureus* osteomyelitis ^59,60^. As cytokine selection was done in U937 and A549, additional cytokines may need to be assessed in SaOS-2 cells. Despite high assay sensitivity, further independent experiments are warranted given the variability observed in the other lines.

### Isolate-specific host cell death induction - TNF-R1 as a gatekeeper of monocyte fate during intracellular S. aureus infection

Manipulating host cell death timing and mode is a key strategy for intracellular escape and bacterial dissemination. Our data confirm cell line- and isolate-specific death rates, signaling pathways, and kinetics consistent with previous reports ^6,9,61-63^. Strikingly, TNF-R1 signaling not only mediates but also modulates U937 cell death, acting as a gatekeeper that shapes the timing and extent of isolate-dependent cytotoxicity, a role far less pronounced in A549 and SaOS-2 cells.

Among the tested cell lines, U937 cells show the fastest and strongest *S. aureus*-induced cytotoxicity, followed by SaOS⍰2 and A549 upon infection with EDCC 5055 or EDCC 5458. SH1000 and EDCC 5464 elicit delayed or minimal responses, affecting only a small proportion of non-professional phagocytes. Flow cytometry indicates distinct death-marker sub-populations, hence fluorescence-based sorting could help distinguish infected from non-infected cells to determine whether caspase negative survivors resist or avoid infection. As reported elsewhere, cytotoxicity is largely restricted to viable bacteria ^64,65^, with moderate effects only from heat-inactivated EDCC 5055.

Initial U937 death requires direct contact or uptake, and involves extrinsic and, for EDCC 5055, also caspase-9-dependent intrinsic apoptosis inducing notable ΔΨm. At 24 hpi, bc-inserts fail to prevent cell death by all isolates except for SH1000, indicating that secreted toxins suffice to drive late⍰stage cytotoxicity, with a potential contribution of elevated toxin levels or nutrient competition, as bacteria remained present under separated conditions. At the same time, apoptotic caspase activity plateaus, whereas isolate dependent caspase-1 activation and 7-AAD staining suggest pyroptotic involvement, though confirmation would require markers such as gasdermin D cleavage or HMGB1 release. Indeed, available studies link several cell death modalities during intracellular *S. aureus* infection with both overlapping and distinct signaling patterns, depending on the infection stage and the host cell types ^39,40,66^. For instance, human neutrophils undergo apoptosis prior to (independently proceeding) lysis ^39^. Conversely, macrophages and epithelial cells that survive initial α-hemolysin-driven inflammatory death exhibit apoptotic features only at later infection stages ^67,68^. Moreover, resource competition within the intracellular niche drives temporal shifts in bacterial and host metabolism, influencing bacterial replication, subcellular localization, and apoptosis of lung epithelial cells ^69^.

Experimental conditions (e.g. assay endpoint, model, and strain choice) strongly shape observed outcomes, and intracellular fates are broadly similar in non⍰phagocytic cells but differ from macrophages ^9^. Consistent with this, viable isolates induce comparable multi-caspase activity in SaOS-2 cells (except EDCC 5464), whereas A549 cells show only marginal caspase activation despite clear detachment. This aligns with reports that pan-caspase inhibitors fail to block cell death and that caspase-independent death can occur even alongside caspase activation ^70,71^.

Remarkably, TNF-R1 knockout in U937 cells eliminates early *S. aureus*-induced cell death, yet promotes broad host cell destruction with pronounced multi-caspase activation and mitochondrial depolarization at later stages, with diminished isolate differences. This supports the notion that intracellular pathogens enhance persistence by suppressing regulated cell death to secure a protected niche ^67,68^. In many cell types, TNF-R1 internalization is required for apoptosis, which can be exploited by pathogens to promote intracellular survival ^27,28,72^.

Interestingly, the isolates used in this study differentially engage or bypass TNF-related signaling in monocytes, suggesting that early, balanced defence against active *S. aureus* relies on TNF-R1-dependent, isolate⍰specific regulated cell death. When TNF-R1 signaling is impaired, prolonged intracellular replication may ensue, ultimately causing delayed and uncontrolled host cell damage, even with otherwise low-cytotoxic isolates such as EDCC 5464.

Consistently, *S. aureus*-infected macrophages are hypothesized to be overwhelmed by high bacterial loads, reducing their bactericidal capacity ^60^, and those that fail to control *S. aureus* are characterized by the formation of intracellular bacterial agglomerations, resulting in cell lysis ^73^. Rodrigues-Lopes et al. describe such a staphylococcal phenotype marked by SpA deficiency, consistent with the absent steady-state SpA expression in all our isolates except the low-cytotoxic EDCC5464. ^9^. Staphylococcal proteins such as SpA and SSL10 directly bind TNF⍰R1, either blocking or triggering host cell death pathways. In epithelial cells and murine models, SpA induces IL⍰8-driven inflammation modulated by TNF⍰R1 shedding ^74-76^, while in osteoblasts it activates NF-κB, apoptosis, and IL-6 release ^77,78^. SSL10-TNF-R1 interaction can promote necroptosis ^79^, although activation of this pathway was not observed in our system. Notably, many cell lines, including A549, lack RIPK3 and are therefore incapable of necroptotic death ^80^.

### Implications for host-pathogen interactions and clinical translation

Anti-inflammatory TNF inhibitors have recently shown benefit in non-bacterial osteomyelitis by attenuating bone loss ^81^. In *S. aureus*-mediated arthritis, ADAM17 inhibition reduces TNF release and has been proposed as an alternative therapeutic approach ^82,83^. ADAM17 also mediates TNF-driven necroptosis in endothelial but not blood cells ^84,85^, while other ADAM proteases (e.g. ADAM9 and ADAM15), upregulated in acute inflammatory lung disease, further modulate TNF-R1 signaling ^86,87^.

In bacterial infections, TNF, TNF-R1, and downstream effectors are essential for clearing intracellular pathogens, and anti-TNF therapy increases infection susceptibility ^88^. Given associated side effects, selective TNF-R1 inhibition has been proposed ^89,90^. Considering our findings, however, delaying regulated phagocyte death may amplify cytotoxicity and inflammation, enabling infected immune cells to act as “Trojan horses” for rapid *S. aureus* dissemination ^4,16-19^. Likewise, only a subset of phagocytes failing to control *S. aureus* suffices to support bacterial replication, escape, and micro-abscess formation ^73^. In mice, TNF-R1 or TNF-R2 deficiency does not alter bacterial burden but differentially shapes neutrophil defence ^91^, while TNF loss impairs phagocytosis, apoptosis, IL-6 release, and macrophage bactericidal activity without affecting granulocytes ^92^.

Considering the pronounced cell-type dependence, targeted inhibition of TNF-R1, TNF-R2, and TNF in additional professional phagocyte and primary cells will be essential to clarify how TNF signaling shapes infection outcomes. Defining the contributions of virulence factors such as SpA and SSL10 will further illuminate underlying mechanisms. Notably, EDCC 5055-induced killing is not restrained even in TNF-R1-expressing U937 cells, and the isolate’s low-inflammatory profile, lacking apoptosis-related pathway enrichment, may reflect bacterial control of this axis. Consistently, blocking autocrine IL-6, TNF-α, or IL-1β secretion in neutrophils abolishes apoptosis protection ^58^. Future work should assess whether *S. aureus* strains modulate TNF-R1 surface levels or internalization, and how pre-stimulation or inhibition of TNF and other cytokines alters these responses.

Although reduced uptake of some isolates, including SH1000, has been linked to cytotoxicity ^6,58^, other studies report no such association ^6,9,10,93^. Consistently, we observe similar intracellular CFU across isolates and between U937_wt_ and U937_ΔTNF-R1_ at 2⍰hpi. Interpretation is limited, however, by inconsistent plating yields caused by bacterial aggregation and non-dividing phenotypes ^7,94^, underscoring the need for parallel high-resolution microscopy. Overall, differential uptake is unlikely to explain the isolate- and cell-type-specific outcomes, but rather differences in intracellular or cytosolic replication capacity.

Phagolysosomal escape and subsequent cytosolic replication are often considered prerequisites for host cell death, especially in non-professional phagocytes ^10,95^, though both processes can occur independently in a strain- and cell-type-specific manner ^6,39^. It requires subcellular manipulation, such as blocking lysosomal acidification or disrupting membranes, driven by agr/saeRS regulators and toxins including α/β-hemolysin, PSMs, or PVL, while other studies highlight roles for systems like graRS or sigB ^3,39,64,68,96-98^. Their contributions vary with experimental conditions, and factors such as leukotoxins/PVL show strong species specificity, predominantly targeting human professional phagocytes ^62,99,100^.

The low-response phenotype of EDCC 5464 likely reflects frameshift mutations in agrC and graS, mobile-element insertions in β-hemolysin, and the absence of additional cytotoxicity-associated genes such as LukED, splABC, and esxB ^37^. Proteomic profiling further shows a lack of PSM and splBDEF expression prior to infection, unlike the other isolates. Notably, esxA can inhibit apoptosis and, together with esxB, is required for bacterial release ^101^; EDCC 5464 retains esxA but lacks esxB, consistent with its persistent, non-cytotoxic, no-escape phenotype. Such behaviour may support prolonged intracellular survival by creating a protected niche. Future work comparing isolate-specific subcellular localization and compartmental markers (e.g. LC3, LAMP-1/-2, acidification) and assessing phagolysosomal escape across cell lines will help clarify the mechanisms governing *S. aureus* intracellular persistence.

Although the isolates share key virulence factors at the genomic and steady-state proteomic level, the absence of secretome quantification during infection remains a major limitation. Defining toxin expression across cell lines will help explain divergent host responses, as varying α-hemolysin levels can trigger distinct death pathways within the same cell type ^70,102^. Consistent with this, Strobel et al. report low, context-dependent cytotoxicity of SH1000, with reduced α-hemolysin, other hemolysins, and PSMs ^6^, which may explain the lack of U937 killing when direct contact was prevented.

## Conclusions

Together, our findings underscore that intracellular *S. aureus* pathogenesis is not a uniform process but emerges from finely tuned interactions between isolate-specific bacterial traits, host-cell type, and TNF-R1-dependent control of balanced monocyte death. Clinical isolates can reach non-canonical, nucleo-proximal localization, distinctly reprogram transcriptional, cytokine, and death pathways, particularly in the professional phagocyte cell lines. By delineating the central role of TNF-R1 in early, regulated monocyte apoptosis, this work provides a framework for understanding infection risks under TNF-targeted therapies and highlights the need to balance suppression of tissue-damaging inflammation with preservation of TNF-R1-mediated antimicrobial functions. In the context of rising antimicrobial resistance, effective treatment of intracellular *S. aureus* will require precisely tailored, tissue- and isolate⍰specific therapeutic strategies.

## Authors contributions

*Concept and project administration:* JF, AW, WSB. *Experiments and data analysis:* AW performed most experiments and data analysis supported by: TB, TG and TS contributed to RNA sequencing and data analysis (Fig. 2 and 3). MJ and GB contributed to SIM imaging (Fig. 1G-J). LFB, OEP and MH contributed to SBF-SEM (Fig. D-F). JB contributed to Fig. 6. LH and SH contributed to TEM (Fig. 1 A-C). NS contributed to Fig. 3E. TZ and UD contributed to proteome analysis. IG contributed to Fig. 1 Av-viii. BK contributed to sequence assembly and annotation. SM, GKM and VA provided reagents. Reagents: SM, GKM, VA *Manuscript writing:* AW, JF, all authors read and approved the manuscript.

## Data and materials availability

All data needed to evaluate the conclusions in this paper are present either in the main text, in the supplementary materials.

## Clinical trial number

not applicable

## Acknowledgement

We thank T. Prenzel and R. Fechter for excellent technical assistance. Library preparation and mRNAseq were performed at the Genomics Core Facility “KFB - Center of Excellence for Fluorescent Bioanalytics” (University of Regensburg, Regensburg, Germany; www.kfb-regensburg.de), thanks to C. Moehle. We thank the Core Unit Fluorescence Imaging and the Department of Biophysics and Biotechnology, University of Würzburg for providing access to the Leica SP8 and Zeiss Elyra 7 SIM.

## List of Abbreviations

ΔΨm: Mitochondrial transmembrane potential
7-AAD: 7-amino-actinomycin D
AWB: Apoptosis Wash Buffer
bc: Boyden chamber
CFU: Colony-forming units
CHX: Cycloheximide
COX1: Cytochrome C oxidase 1
DEG: Differentially expressed gene
FASP: filter-aided sample preparation
GO:BP: Gene onotology biological processes
hpi: Hours post infection
LAMP: Lysosome-associated membrane protein
LB: Luria-Bertani broth
Log2FC: Log2-transformed fold change
MAPK: Mitogen-activated protein kinases
Min_adj_score: Minimal adjusted enrichment score
MOI: Multiplicity of infection
nCI: Normalized cell index
NF-κB: Nuclear factor κB
OD_600_: Optical density at 600 nm
Padj: Adjusted p-value
PARP-1: Poly(ADP)ribose-polymerase 1
PCA: Principal component analysis
PI: Propidium iodide
PTMs: Post-translational modifications
*S. aureus*: *Staphylococcus aureus*
SBF-SEM: Serial block face scanning electron Microscopy
SIM: Structured Illumination Microscopy
TEM: Transmission electron micrsocopy
TNF: Tumor necrosis factor α
TNF-R1/-R2: TNF-receptor 1/ -2
zVAD.fmk: Benzyloxycarbonyl-Val-Ala-Asp(OMe)-fluoromethylketone [Pan-caspase inhibitor]

## eMail contact

Walter, Annika

Beck, Johanna

Bischler, Thorsten

Jungblut, Marvin

Breitsprecher, Leonhard F.

Schaefer, Nicole

Hofmann, Lucia

Ziesmann, Tanja

Haerteis, Silke

Gadjalova, Iana

Distler, Ute

Beliu, Gerti

Psathaki, Olympia Ekaterini

Hensel, Michael

Schneider-Brachert, Wulf

Graefenhan, Tom

Stempfl, Thomas

Kieninger, Baerbel

Mühlen, Sabrina

Alt, Volker

Mannala, Gopala-Krishna

Fritsch, Juergen

