## Supplemental figures for "*Staphylococcus aureus* triggers isolate-specific host transcriptional responses alongside TNF-R1 regulated cell death"

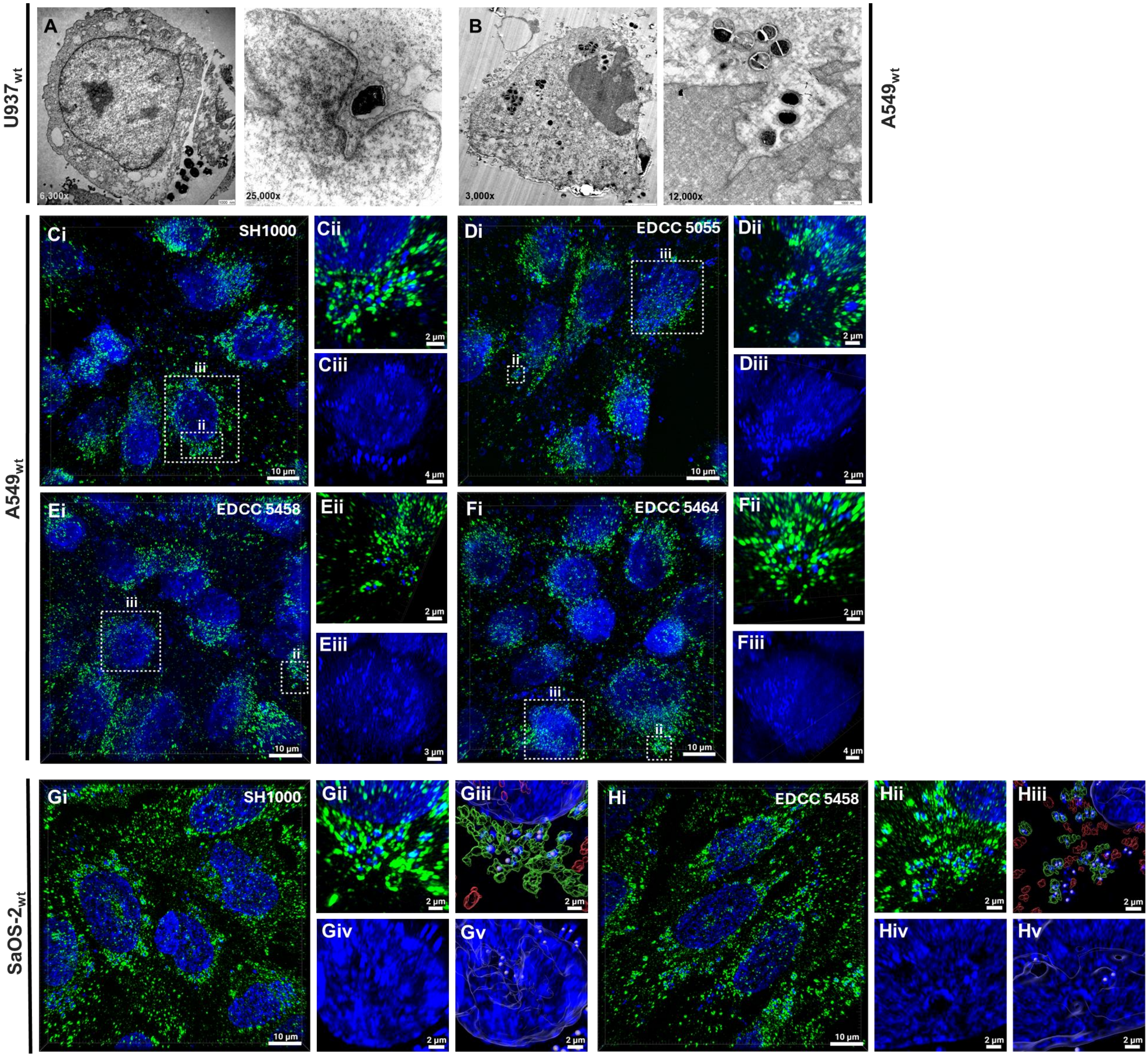


**Fig. S3 Peri- and intranuclear, LAMP-1 associated localization of *S. aureus* at 2 hpi and MOI 100.** Each cell line was infected with *S. aureus* at MOI 100 using a gentamicin protection assay and cells were fixed at 2 hpi. Transmission electron microscopy of peri- and intranuclear localization (**A**) EDCC 5055 in U937 and (**B**) EDCC 5464 in A549 cells. 3D-rendering of (**C-F**) A549 and Maximum intensity projection of (**G-H**) SaOS-2 cells infected with *S. aureus* (**C, G**) SH1000, (**D**) EDCC 5055, (**E**, **H**) EDCC 5458, and (**F**) EDCC 5464 imaged by SIM. DNA was stained by DAPI (blue) and LAMP-1 was stained by mouse anti-LAMP1 and AF488 (green). (**Cii/iii – Fii/iii)** Zoom-in of boxed region in Ci-Fi. (**Gii/iii, Hii/iii**) Surfaces and spots generated after image analysis in Imaris (11.0.0). Surfaces defining the nucleus are visualized by a blue outline, spots corresponding to bacterial DNA are highlighted by white spheres. Surfaces generated around LAMP1 signal are classified based on distance to the closest bacterial spot. Surfaces closer than 0.25 µm coloured in green, while surfaces at a greater distance are coloured in red. (**Giv, Hiv**) DAPI stained nucleus of a cell infected with EDCC 5055/5464 (not depicted in Gi). Strong DAPI staining consistent with bacterial DNA stain can be observed inside the nucleus or located within an invagination of the nuclear lamina. (**Gv**, **Hv**). Spots are identified inside the generated nucleus surface.

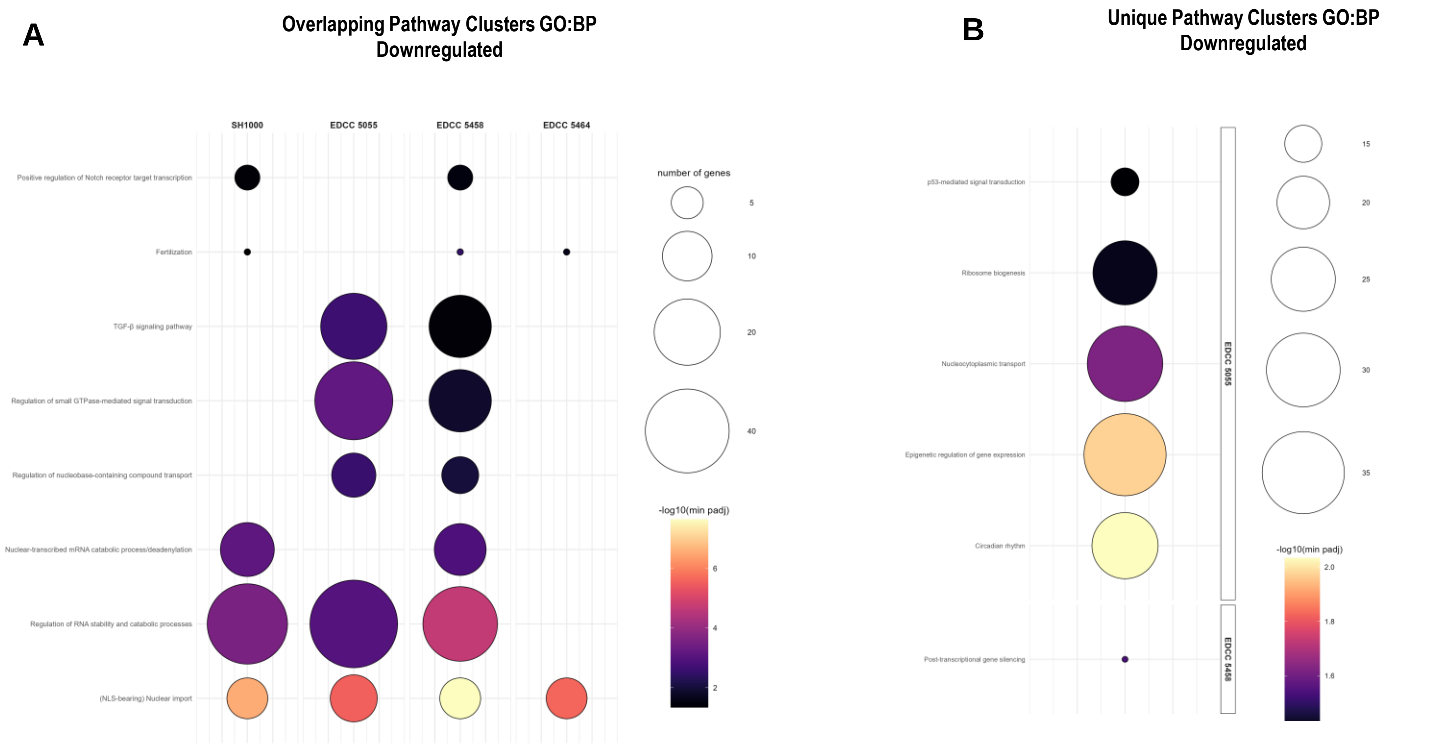


**Fig. S10 Intracellular *S. aureus* infection of U937 cells induces distinct downregulated pathway clusters dependent on each isolate.** Significantly (padj≤0.05), at least two-fold upregulated protein-coding genes were analyzed for enriched pathways (Gene Ontology Biological Processes) using g:Profiler ordered query, and clustered based on Jaccard similarity of gene sets, using hierarchical clustering (average linkage) and a fixed cut height of 0.5. The adjusted enrichment score of a driver pathway (minimal padj) is shown representatively for each cluster (-log10(min_adj_score)) and bubble size indicates gene set size, for (**A**) overlapping and (**B**) unique pathway clusters of *S. aureus* isolates compared to untreated controls.

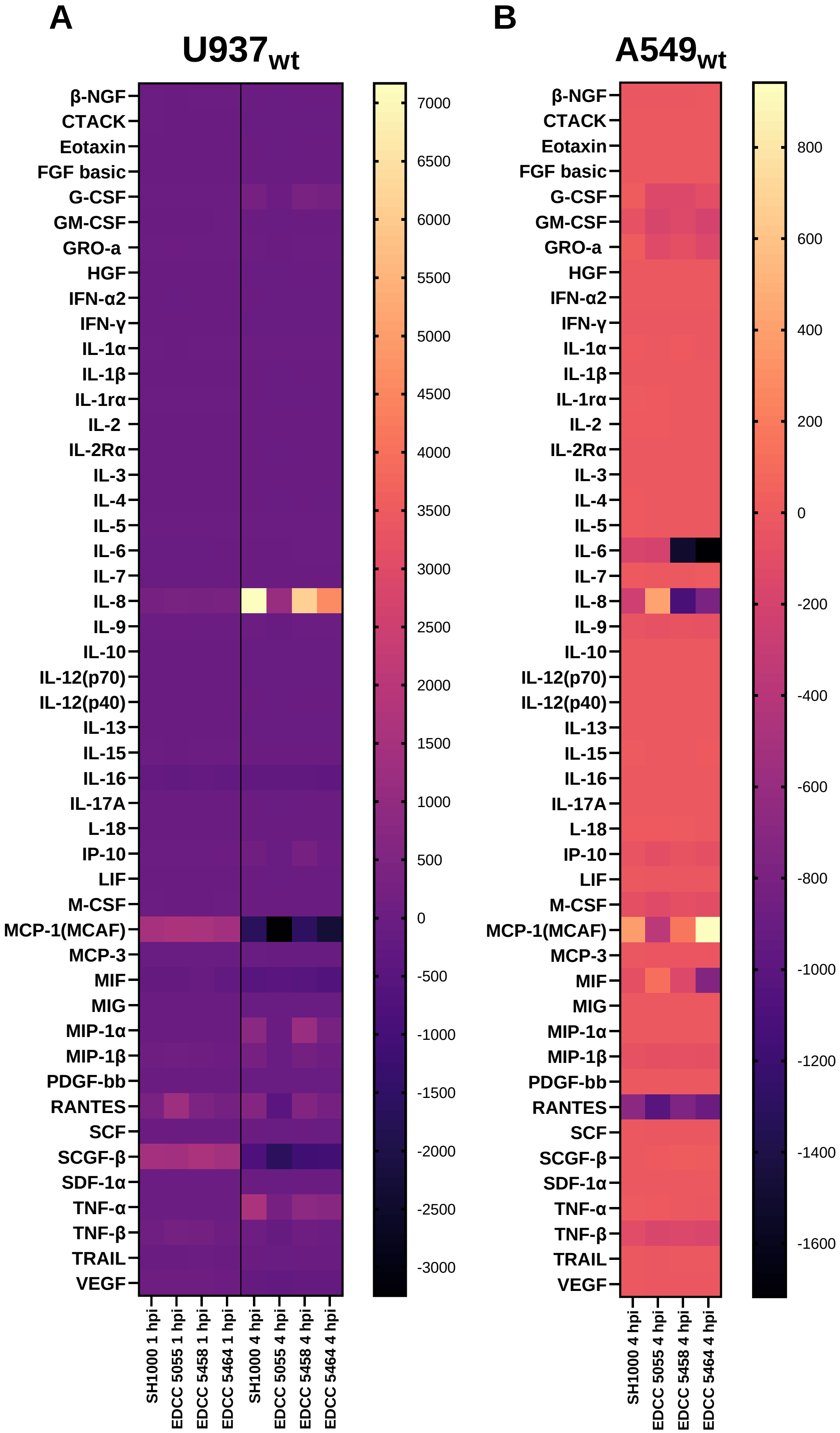


**Fig. S12 Luminex 48-plex ELISA of *S. aureus*-infected (A) U937wt at 1 hpi and 4 hpi and (B) A549wt cells at 4 hpi (n=1), MOI 30.** Fluorescence intensities normalized to uninfected control cells and background.

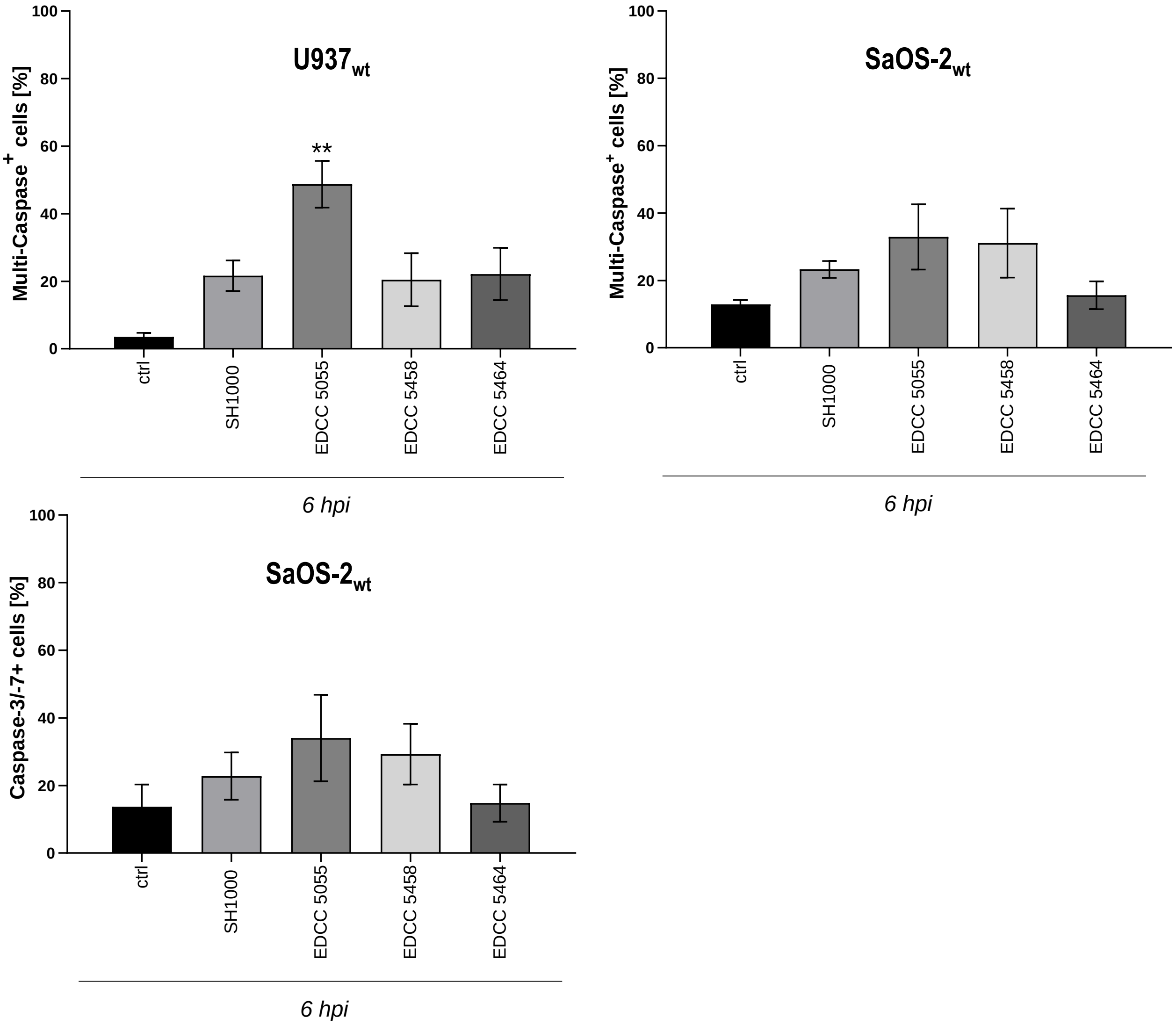


**Fig. S13 Flow cytometry analysis of activity of multiple caspases in (A) U937wt and (B) SaOS-2 cells and (C) caspase-3/-7 in SaOS-2 cells upon *S. aureus* infection at 6 hpi following a gentamicin protection assay.** Data show mean ± SEM of n≥3 independent experiments. Statistical analysis was performed using One-way ANOVA with Tukey's Multiple Comparisons comparing means between all conditions. Signifance is shown for comparison against uninfected controls. **p<0.01.
